# The genomes of invasive coral *Tubastraea* spp. (Dendrophylliidae) as tool for the development of biotechnological solutions

**DOI:** 10.1101/2020.04.24.060574

**Authors:** Giordano Bruno Soares-Souza, Danielle Amaral, Daniela Batista, André Q. Torres, Anna Carolini Silva Serra, Marcela Uliano-Silva, Luciana Leomil, Aryane Camos Reis, Elyabe Monteiro de Matos, Emiliano Calderon, Vriko Yu, Francesco Dondero, Saulo Marçal de Sousa, David Baker, Aline Dumaresq, Mauro F. Rebelo

**Affiliations:** Bio Bureau Biotechnology, Rio de Janeiro, RJ, Brazil; Institute Senai of Innovation in Biosynthetics, SENAI CETIQT, Rio de Janeiro, RJ, Brazil; Biophysics Institute Carlos Chagas Filho, Federal University of Rio de Janeiro, RJ, Brazil; Berlin Center for Genomics in Biodiversity Research (BeGenDiv), Berlin, Germany; Laboratory of Genetics and Biotechnology, Federal University of de Juiz de Fora, Juiz de Fora, MG, Brazil; Institute of Biodiversity and Sustainability (NUPEM), Federal University of Rio de Janeiro, RJ, Brazil; Coral Vivo Institute, BA, Brazil; The Swire Institute of Marine Science and School of Biological Sciences, The University of Hong Kong, Hong Kong, SAR; Università degli Studi del Piemonte Orientale (UNIPO)

**Keywords:** Sun Coral, gene-environment, Brazilian coast

## Abstract

Corals have been attracting huge attention due to the impact of climate change and ocean acidification on reef formation and resilience. Nevertheless, some species like *Tubastraea coccinea* and *T. tagusensis* have been spreading very fast replacing the native ones which affect the local environment and decrease biodiversity of corals and other organisms associated with them. Despite some focal efforts to understand the biology of these organisms, they remain understudied at the molecular level. This knowledge gap hinders the development of cost-effective strategies for both conservation and management of invasive species. In this circumstance, it is expected that genome sequencing would provide powerful insights that could lead to better strategies for prevention, management, and control of this and other invasive species. Here, we present three genomes of *Tubastraea* spp. in one of the most comprehensive biological studies of corals, that includes flow cytometry, karyotyping, transcriptomics, genomics, and phylogeny. The genome of *T. tagusensis* is organized in 23 chromosomes pairs and has 1.1 Gb, the *T. coccinea* genome is organized in 22 chromosome pairs and has 806 Mb, and the *Tubastraea* sp. genome is organized in 21 chromosome pairs and has 795 Mb. The hybrid assembly of *T. tagusensis* using short and long-reads has a N50 of 227,978 bp, 7,996 contigs and high completeness estimated as 91.6% of BUSCO complete genes, of *T. coccinea* has a N50 of 66,396 bp, 17,214 contigs and 88.1% of completeness, and of *Tubastraea* sp. has a N50 of 82,672 bp, 12,922 contigs and also 88.1% of completeness. We inferred that almost half of the genome consists of repetitive elements, mostly interspersed repeats. We provide evidence for exclusive Scleractinia and *Tubastraea* gene content related to adhesion and immunity. The *Tubastraea* spp. genomes are a fundamental study which promises to provide insights not only about the genetic basis for the extreme invasiveness of this particular coral genus, but to understand the adaptation flaws of some reef corals in the face of anthropic-induced environmental disturbances. We expect the data generated in this study will foster the development of efficient technologies for the management of coral species, whether invasive or threatened.

## Sequencing the genome is a landmark in the history of a species

Corals are among the planet’s most stunningly beautiful organisms and they exist in an amazing variety of shapes, sizes and colors. Corals reproduce sexually or asexually ^1^, live alone or in colonies, with or without symbionts ^2^. Corals form blooming reefs in ecosystems that range from low productivity shallow hot waters to nutrient-rich banks in the coldest depths of the ocean floor ^3^.

As enchanting as they are, very little is known about these amazing creatures at the molecular level. To date, only seven coral species have had their genomes deposited in the NCBI Genome database (see Box 1 and Table 1).

### Box 1

**What do we know about coral genes thought to confer invasiveness?**

First, genomes of corals are a rarity. Of 9188 Eukaryotes genome assemblies deposited in NCBI Genome Database, just 29 are from Cnidaria and only eight are from Scleractinia order (stony corals) species (Table 1). Even in specialized databases, such as the Reef Genomics (reefgenomics.org) and the University of Kiel’s Comparative genomics platform from (Compgen from Bosch’s lab - http://www.compagen.org), the number of available studies is modest, sixteen projects in the former database and five datasets in the latter.

Regarding the genes involved in coral development, we know that some genes expressed in bilateria clade, such as those from *Wnt* gene family, are present in Cnidarian genomes and are expressed in early life stages of Scleractinian corals ^4^

We know that *T. tagusensis* is able to trigger mouth regeneration and regenerate a new polyp faster than *T. coccinea* ^5^, but we know nothing about genes involved in these processes. A Hemerythrin-like protein was found expressed only in the regeneration process, suggesting that it is a “regeneration-specific” gene. A few studies have found genes related to differentiation and regeneration in hydra and salamander ^6–9^ that were up-regulated in the sea anemone *N. vectensis* during the regeneration process.

The calcification of the skeletal organic matrix is thought to be related to the expansion of Carbonic anhydrases (CAs) genes in hard corals and constitutes the main difference from Corallimorpharians ^10,11^.

We know that *Tubastraea* spp. can and do reproduce sexually ^12^ but very little is known about the genes involved in triggering this process. A gene called *euphy*, discovered in *Euphyllia ancora*, has been found to be overexpressed before the reproductive season in ovarian somatic cells and accumulates as a yolk protein, essential to embryonic development ^13^. Two other yolk proteins were identified in corals, vitellogenin and egg protein, which are produced in *E. ancora* and it is suggested that vitellogenin is also present in other Cnidaria ^14^.

**Table 1:** Statistics overview from our haploid draft assembly compared to other coral genomes available in <u>NCBI’s Genbank</u>. The *Tubastraea* spp.

| Species | Genome Size (Mb) | Contig N50 (bp) | Contigs | GC% | Complete BUSCO | Reference |
| --- | --- | --- | --- | --- | --- | --- |
| *Tubastraea tagusensis* | 1,125 | 227,044 | 7,996 | 38.92% | 91.6% | This study |
| *Tubastraea coccinea* | 807 | 66,396 | 17,214 | 38.76% | 88.1% | This study |
| *Tubastraea* sp. | 795 | 82,672 | 12,922 | 38.87% | 88.1% | This study |
| *Acropora digitifera* | 447 | 10,915 | 54,401 | 40.5% | 52.6% | <sup>33</sup> |
| *Acropora millepora* | 386 | 36,677 | 20,440 | 35.1% | 90.5% | <sup>34</sup> |
| *Montipora capitata* | 615 | 24,266 | 50,174 | 39.6% | 81% | <sup>35</sup> |
| *Orbicella faveolata* (v1) | 48 | 3,840 | 155,027 | 41.9% | 81.1% | <sup>36</sup> |
| *Orbicella faveolata* (v2) | 486 | 12,468 | 55,729 | - | 85.2% |  |
| *Pocillopora damicornis* | 234 | 25,941 | 53,034 | - | 88.1% | <sup>37</sup> |
| *Porites rus* | 470 | 5,139 | 81,422 | 38.8% | 72.5% | <sup>38</sup> |
| *Stylophora pistillata* | 400 | 20,604 | 37,615 | 39.7% | 88% | <sup>39</sup> |

This lack of knowledge hinders the development of strategies that control specifically invasive species like *Tubastraea* and do not harm the ecosystem. The impact of climate change on coral reefs has dominated the research agenda while invasive coral species remain understudied, even in light of their significant role in the loss of biodiversity in rocky shores.

*Tubastraea* (Cnidaria: Dendrophylliidae) is a fast growth ^15^ azooxanthellate coral with no significant substrate specificity ^16^, with ability to produce planulae both sexually and asexually ^17,18^ as well as to fully regenerate from undifferentiated coral tissue ^5^. Since the first reports in the late 1930s, in Puerto Rico and Curacao ^19,20^, *Tubastraea* spp. – coral species native to the Indo-Pacific Ocean ^21^ – have spread rapidly throughout the Western Atlantic Ocean. Invasive *Tubastraea* corals are found in the Caribbean Sea, the Gulf of Mexico and have been detected discontinuously along 3,850 km of the coast of Brazil (from 2°30’S to 26°30’S) ^20,22–28^, occupying up to 95% of the available substrate in some regions ^12^. Recently, *Tubastraea* corals were found around Eastern Atlantic islands including the Canary Islands ^29,30^.

Without innovation in control methods, dispersion is expected to continue, as desiccation in drydocks and physical removal cannot be applied in a timely and cost-effective manner, or risks inadvertently contributing to further dispersion.

We know that gene-environment interactions often result in gene expression changes. Thus, characterization of the *Tubastraea* spp. genomes should help to elucidate the molecular mechanisms of tolerance, resistance, susceptibility and homeostasis that could lead to better conservation strategies for corals as well as specific methods of control for this invasive species. Also, the assemblies of the complete mitochondrial genome of three morphotypes of *Tubastraea* whose complete genomes were sequenced, together with three mitochondrial markers sequenced to more eighteen specimens, allowed us to better understand the occurrence of other *Tubastraea* species occurring at Brazilian coast.

Here, we present the draft genomes of *Tubastraea* spp., assembled using short and long reads, aggregated with RNA-seq data, flow cytometry and karyotype information and morphological characterization of the colonies. In comparison to other corals, they are the largest genomes sequenced to date, and we made one of the most comprehensive efforts to elucidate genomic organization in a coral genus.

## We sequenced three morphotypes of *Tubastraea* colonies from southeast Brazil

After collection in Angra dos Reis (23°3.229’S; 44°19.058’W) and Arraial do Cabo (22o57’56”S; 41o59’36”W) - SISBIO collection authorization number 68262 - three distinct *Tubastraea* colonies were maintained in our laboratory. Two species reported for the Brazilian coast were sampled, *Tubastraea tagusensis* Wells, 1982 and *Tubastraea coccinea* Lesson, 1830 ^24^. Of the two colonies sampled in Angra dos Reis, *T. tagusensis* morphotype (SC065) has a corallum phaceloid forming a bush-shaped colony and elliptical corallites which are budding from a broad base and project up to 40 mm above the coenosteum (Figure 1 A-C). Despite recent concerns raised by ^31^, we maintain this morphotype temporarily identified as *T. tagusensis. Tubastraea coccinea* morphotype (SC082) has a corallum placoid forming spherical shape colony and circular corallites originating from the same basal coenosteum, with closely-spaced among them, and projecting up to 6 mm above the coenosteum (Figure 1 G-I). The third colony (SC100) collected in Arraial do Cabo is a morphological variation of *T. coccinea* that was recently identified as *Tubastraea aurea* by ^31^. It has a corallum placoid and corallites originate from the same basal coenosteum budding moderate widely-spaced among mature corallites (Figure 1 D-F), which project up to 10 mm above the coenosteum (Figure 1 D-F). Some corallites present part of the cycle septal with the initial stage of the Pourtalés Plan.

**Figure 1:**
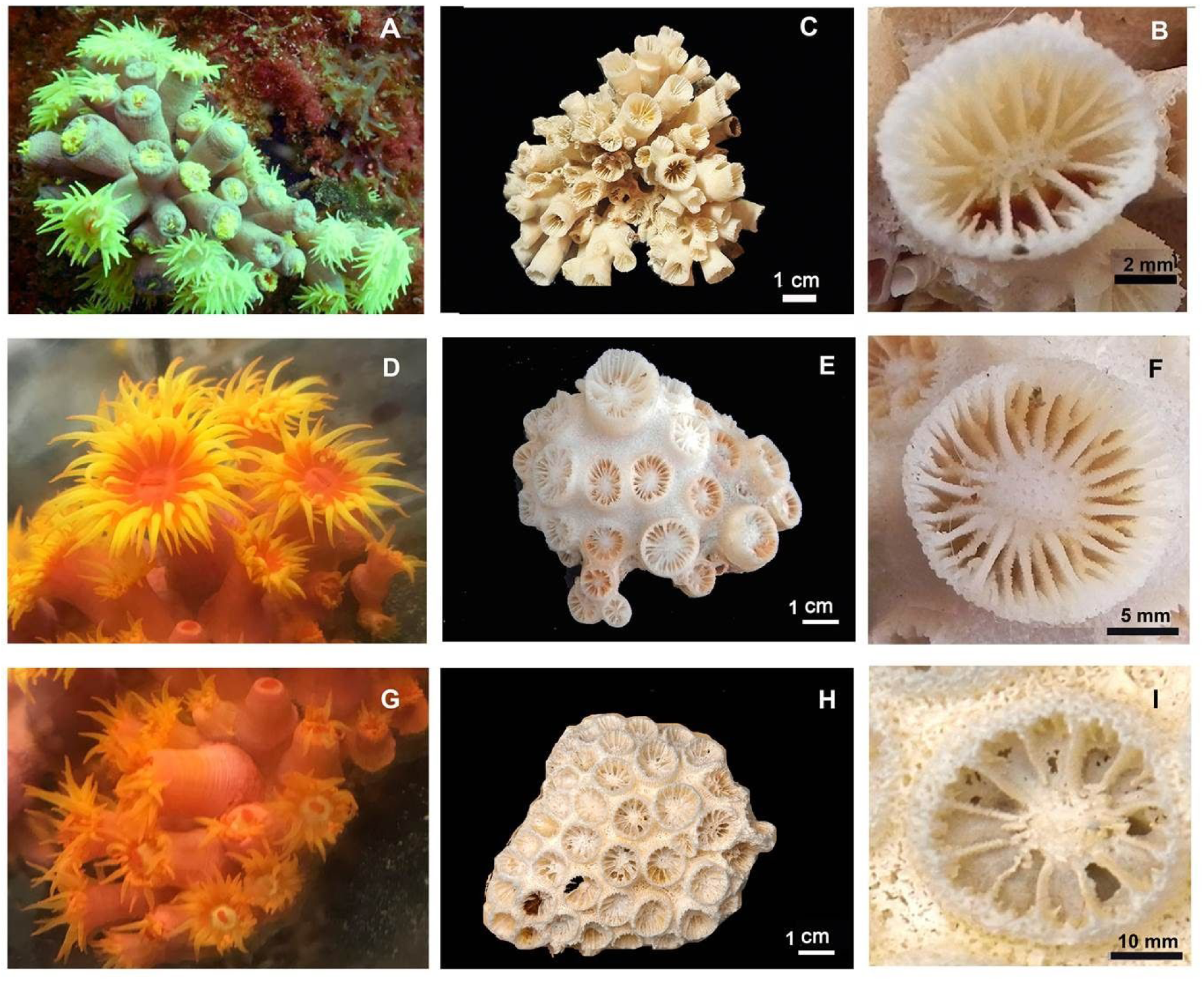
*In vivo* colonies and details of their skeletons: (A) Morphotype of *T. tagusensis* (SC065) *in viv*o, with yellow polyps connected by coenosarc with the same colour; (C) Corallum phaceloid forming colony with 9.1 cm in diameter; (B) Detail of septa arrangement, with primaries (S1) and secondaries (S2) septa reaching the center of corallite. (D) Morphotype of *Tubastraea* sp. (SC100) *in vivo*, with coenosarc orange-red in color, while tentacles and mouth are yellow and red-orange bright, respectively; (E) Corallum placoid forming colony with 6.7 cm in diameter; (F) Detail of septa arrangement, with part of cycle septal with initial stage of Pourtalés Plan. (G) Morphotype of *T. coccinea* (SC082) *in vivo*; (H) Corallum placoid forming colony with 6.7 cm in diameter; (I) Detail of septa arrangement, with S1 and S2 reaching the center of corallite.

All nucleic acid extractions were performed in clone polyps. Colonies morphology were analyzed and the skeletons were preserved in order to be deposited in the zoological collection of the National Museum of Brazil.

The mitochondrial genome of the three morphotypes of *Tubastraea* whose genomes were sequenced in this work have more than 99% of identity with NC_030352.1 and KX024566 (*T. tagusensis* and *T. coccinea*, Capel and collaborators^32^), Indeed, *T. tagusensis* (SC065), *T. coccinea* (SC082), *Tubastraea* sp. (SC100) and the two genomes described by Capel and collaborators^32^, clustered in a high supported monophyletic clade (highlighted with a black star; Figure 2A). Inside of that, DNA and RNA assemblies of SC065 grouped together with NC_030352.1 in a subclade with 100% of bootstrap (clade with a red star; Figure 2A), while both assemblies of SC082 and the DNA assembly of SC100 get together in another subclade with KX024566 (clade with a green star; 100% of bootstrap; Figure 2A). Although SC100 has been grouped with *T. coccinea*, it is in a basal position in this subclade and has a different pattern of polymorphisms in comparison with the other *Tubastraea*. Of the 49 sites, 31 are shared with *T. coccinea*, eleven with *T. tagusensis* and 7 are specific of this morphotype (Figure 2A). Like described by Capel and colleagues ^32^, NC_026025.1 grouped with *Dendrophyllia arbuscula* (100% bootstrap) out of the *Tubastraea* clade.

**Figure 2.**
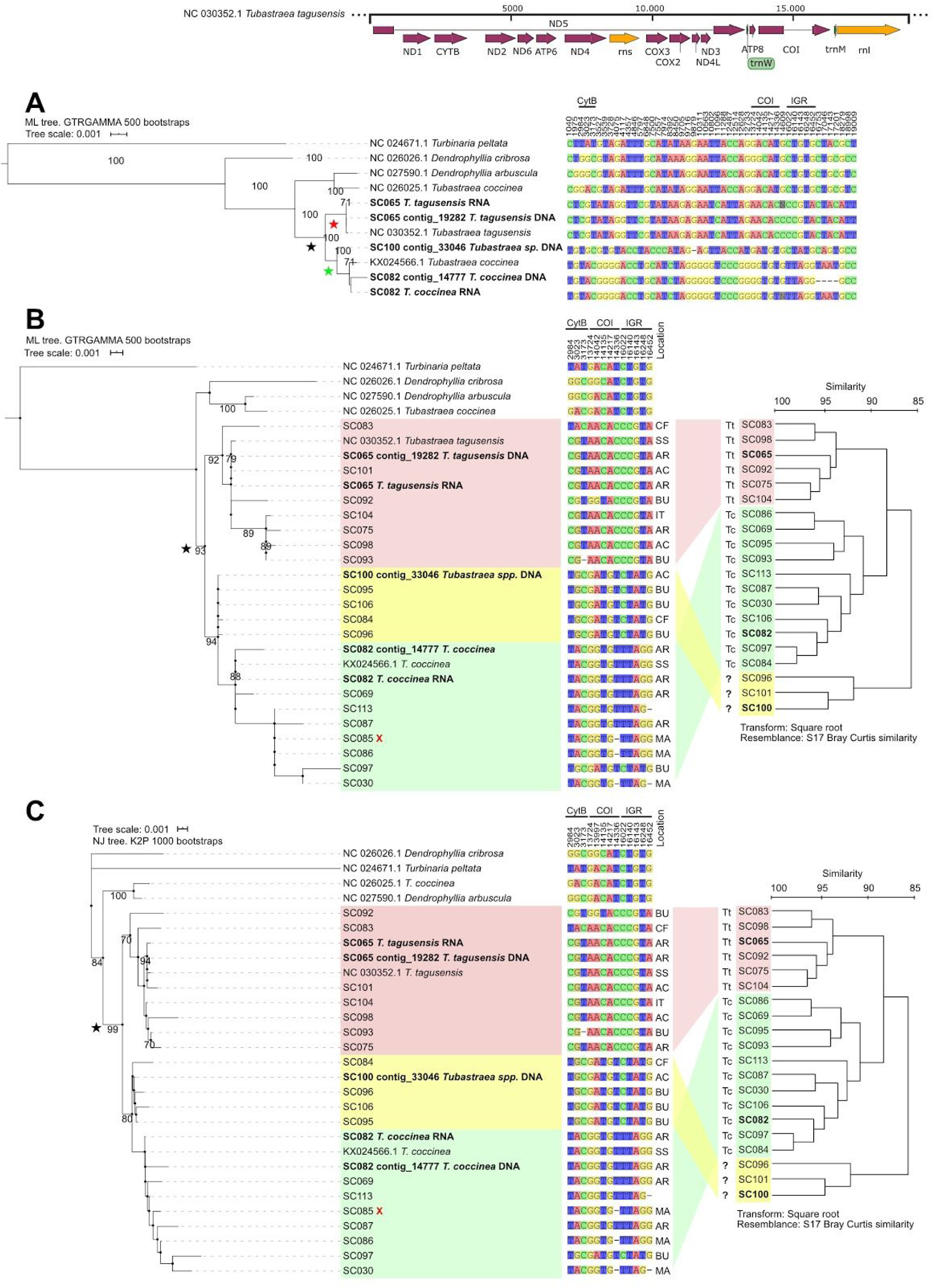
Mitochondrial analysis and comparison of the molecular and morphology identification. A - ML tree using the whole mitochondrial genome of SC065, SC082, SC100 and other Dendrophylliidae. *Tubastraea* clade is highlighted with a black star and *T. tagusensis* and *T. coccinea* subclades with a red and green ones. All forty-nine polymorphic sites occurring in between the three morphotypes sequenced in this work are shown. The schematic diagram of the mitochondrial genome indicates where the positions taking NC_030352.1 as reference are. B - ML tree built with the sequences of the CytB, COI and IGR of eighteen specimens (“SC” prefix) sampled in Buzios (BU), Cabo Frio (CF), Angra dos Reis (AR), Arraial do Cabo (AR), Ilhas Tijuca (IT) and Macae (MA). NC_030352.1 and KX024566 were collected in São Sebastião (SS). The red “X” in SC085 indicates that this morpho-type could not be identified through their morphological characters. For a better visualization, the tree based on morphological characters was mirrored. *T. tagusensis* (Tt) clade is coloured in red, *Tubastraea sp.* (?) - yellow and *T. coccinea* (Tc) - green. Thirteen positions occuring in the three gene markers are shown. C - NJ tree built with the same sequences used in the analysis shown on figure B. Only bootstraps higher than 70% were shown in the trees.

**Figure 3:**
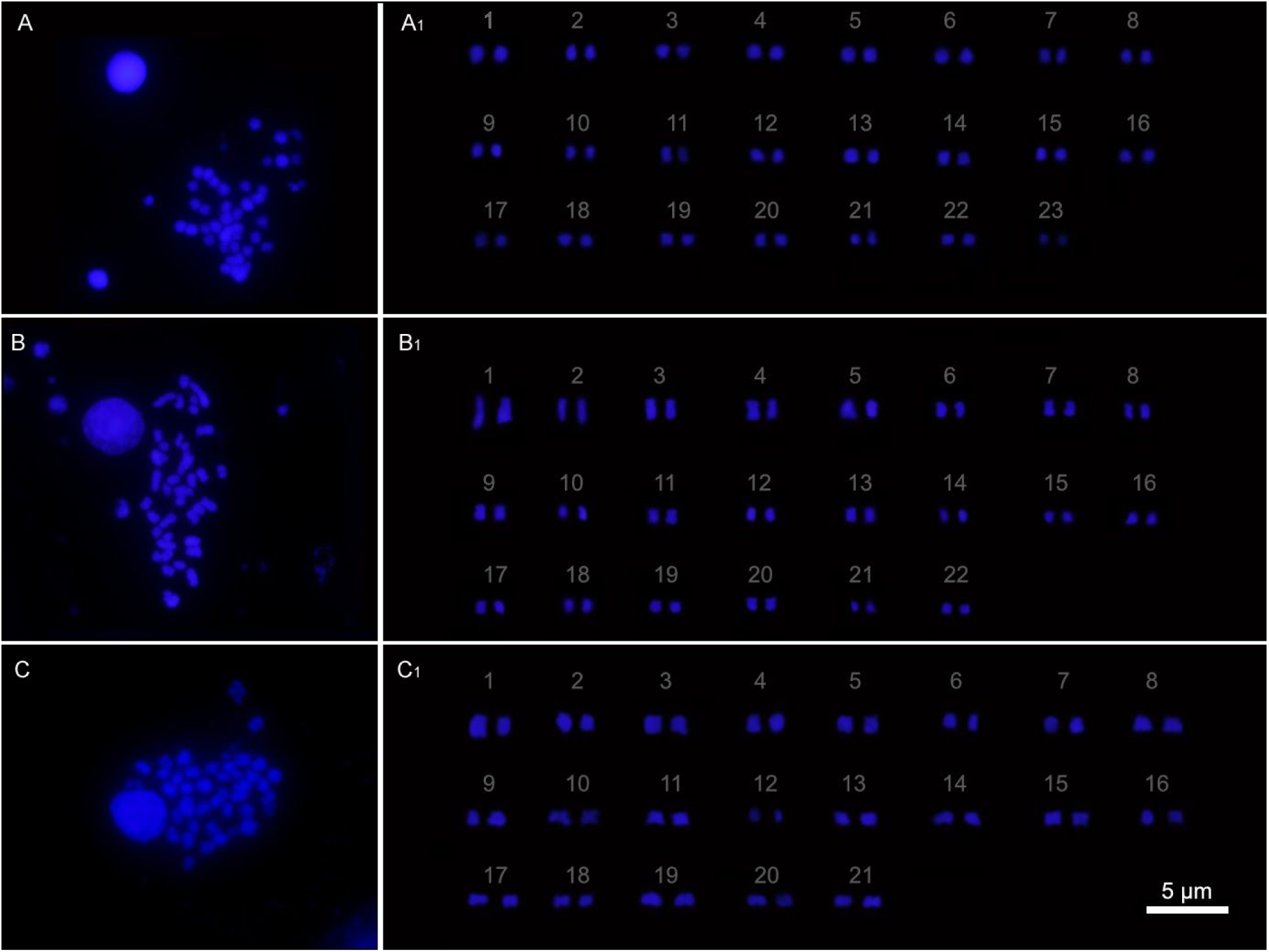
Representative metaphase and karyogram of (A and A1) *T. tagusensis* (2n= 46), (B and B1) *T. coccinea* (2n= 44), and (C and C1) *Tubastraea* sp. (2n= 42). Two metaphase plates of each morphotype were counted. Scale bar 5 µm.

**Figure 4.**
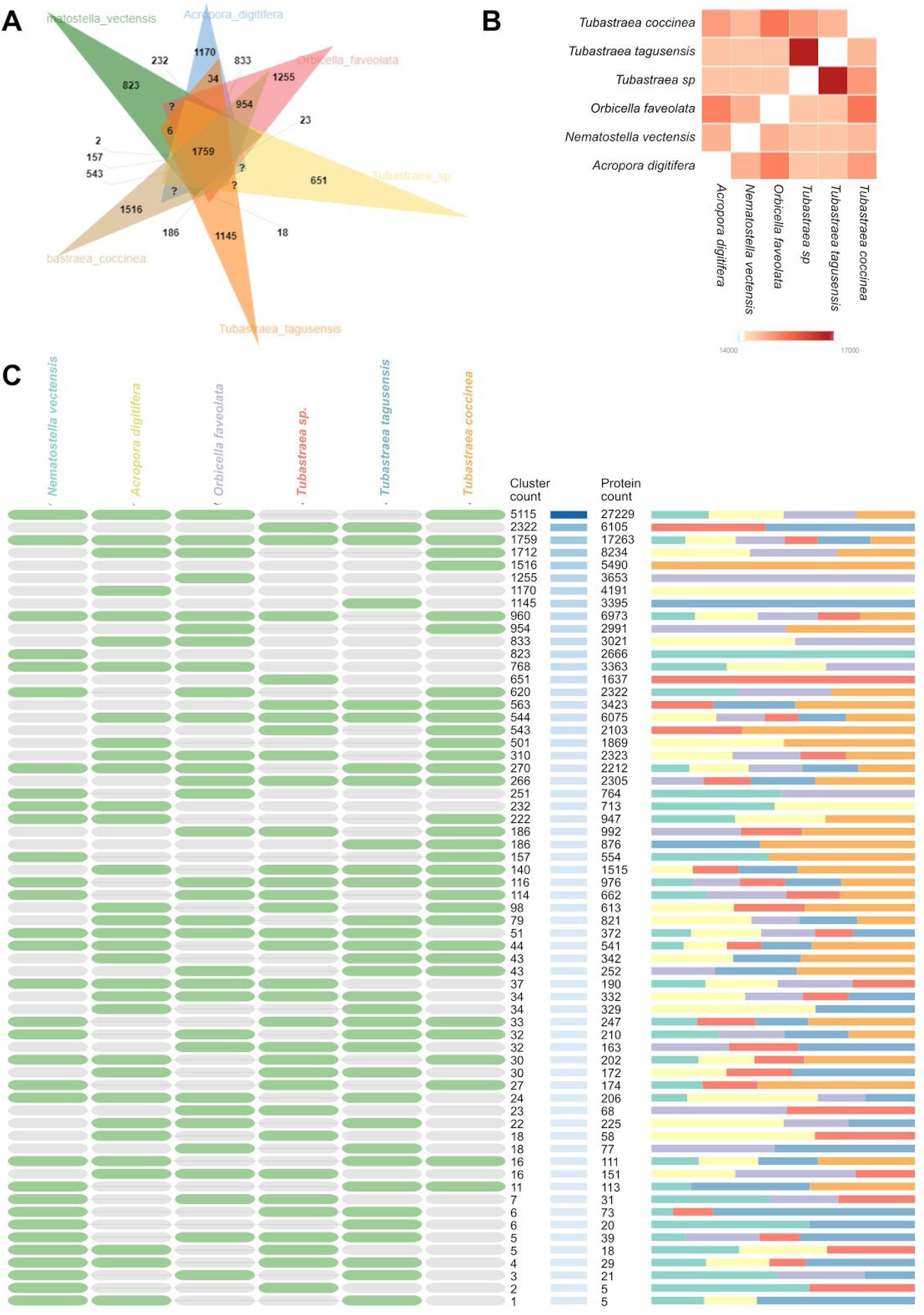
Whole-genome orthologous comparisons across 6 Cnidaria species: *Nematostella vectensis, Acropora digitifera, Orbicella faveolata, Tubastraea coccinea, Tubastraea tagusenis* and *Tubastraea* sp. **(A)** Venn diagram showing the distribution of shared gene families among Cnidaria species. **(B)** Pairwise overlapping cluster numbers heatmap, darker red shows increased cluster sharing between species. **(C)** Overlapping orthologous gene clusters across Cnidaria species.

Beside the whole mitochondrial genome analysis, we investigated the phylogenetic relationship and polymorphisms occurring in three gene markers: cytochrome oxidase I (COI), cytochrome oxidase b (CytB), and a intergenic spacer (IGR), of other seventeen specimens of coral from Rio de Janeiro and one with unknown location (SC113). The phylogenetic trees using the concatenated alignment of these regions show that all specimens grouped in a monophyletic clade in both maximum likelihood (ML) and neighbor joining (NJ) trees (clades with a black star and 93% and 99% of bootstrap; Figure 2B and 2C). Six specimens morphologically identified as *T. tagusensis* (SC075, SC065, SC083, SC092, SC098 and SC104), one *T. coccinea* (SC093) and one unidentified (SC101) get together with NC_030352.1 (*T. tagusensis*) in a monophyletic clade (92% and 70% of bootstrap; red clades in figure 2B and 2C). Indeed, they have the same nucleotide in all polymorphic sites occurring in these regions, except for SC083 (T:2984, A:3023 and C:3173) and SC092 (G:13724, G:13997 and T:14136) that have three divergent sites and SC093 that has a deletion at the position 3173 because of the lack of the beginning of the COI marker (figure 2B and 2C).

Eight specimens identified as *T. coccinea* (SC030, SC069, SC082, SC084, SC086, SC087, SC095, SC097, SC106, SC113), one as *T. tagusensis* (SC085), and two unidentified (SC096 and SC100) clustered with KX024566 (*T. coccinea*) in a high supported monophyletic clades (94% and 80% of bootstrap in ML and NJ trees; figure 2B and 2C). In this clade, four polymorphic sites with the same nucleotide of *T. tagusensis*, which split it into two monophyletic groups (yellow and green clades), although with no statistical support. Taking NC_030352.1 as reference, they are at the positions 3023 (G/A), 14042 (A/G), 16022 (C/T), 16248 (T/G) considering the yellow and green clades, respectively (Figure 2B and 2C).

## *Tubastraea* spp. have large genomes and different number of chromosome pairs

Through flow cytometry analysis, we estimate that the size of the haploid genome of *T. tagusensis* and *Tubastraea* sp. is 1.3 gigabases (Gb), while *T. coccinea* has 1.1 Gb (Supplementary table 2).

The representative karyogram was assembled from two metaphases plates of each specimen, using genetic material from a pool of planulae from colonies of the same morphotypes that have had its genome sequenced. Planulae were treated with 0.1% colchicine and metaphases plates were prepared. The three morphotypes show diploid karyotypes and a different number of chromosomes. *T. tagusensis* has 46, *T. coccinea*, 44 and *Tubastraea* sp., 42 (Figure 2). The Ideograms (Supplementary Figure 4) shows the medium size of the short and long arms of the chromosomes. To our knowledge, this is the first Dendrophylliidae genomic study to present evidence about the number of chromosomes and ploidy.

DNA extraction of *T. tagusensis* using the *DNeasy* Blood & Tissue kit yielded a DNA of about 10 kb in size with a DNA Integrity Number (DIN) of 6.4 (Supplementary Figure 5), free of proteins, and it was used to construct the Illumina library. DNA extraction using the CTAB buffer protocol yielded a DNA of about 23 kb in size and DIN = 8.0 (Supplementary Figure 6) and also free of proteins and other contaminants. It was used to construct all the SMRTbell libraries and Illumina libraries from *T. coccinea* and *Tubastraea* sp.. The DNA extraction from *T. coccinea* yielded a DNA > 60 kb in size and DIN = 8.0 (Supplementary Figure 7). And the DNA extraction from *Tubastraea* sp. yielded a DNA around 14 kb in size (Supplementary Figure 8).

## We recovered more than 80% of BUSCO complete genes in the transcriptome

The stranded paired-end RNA sequencing raw data consisted of about 60 million reads for

*T. coccinea* and *T. tagusensis* and 30 million reads for *Tubastraea* sp.. RNA raw data was quality trimmed and filtered yielding about 7 billion bases for *T. tagusensis*, 6 billion for *T. coccinea* and 3.8 billion for *Tubastraea* sp.

Trinity assembled a *de novo* transcriptome with a little more than 350,000 transcripts for *T. tagusensis* and *T. coccinea* and 200,000 for *Tubastraea* sp. with an average length ranging from 600 bases for *T. coccinea* and 800 for *T. tagusensis* and *Tubastrea* sp.. Transcriptome assembly N50 ranged from about 1,400 kb for *T. tagusensis* and *Tubastrea* sp. to 850 for *T. coccinea*. Of these reads, more than 90% mapped to the transcriptome and 75% mapped to the genome for each morphotype. We recovered from 80 to 98% of complete genes in a search for orthologous genes using the BUSCO metazoa database and retrieved more than 4,000 (about 30-40%) near-complete protein-coding genes with reads mapping to more than 80% of the estimated protein length. The final set consisting more than 100 thousand transcripts included only sequences with coding and homology evidence (Table 1).

## We recovered more than 88% of BUSCO complete genes in the genomes

A total of 383 Gb of paired-end DNA data were obtained of *T. tagusensins* using an Illumina HiSeq X. Of *T. coccinea* and *Tubastraea* sp., 152 Gb and 89 Gb were obtained using a Illumina NextSeq550, respectively. After sequencing on a PacBio Sequel I platform 92 Gb of sub-reads raw data were generated for *T. tagusensis*, 157 Gb for *T. coccinea*, and 135 Gb for *Tubastraea* sp.

Short-reads were quality trimmed and filtered retaining about 85% of both reads and bases. Using MaSuRCA assembler and Purge Haplotigs software to filter allelic contigs, we estimated the haploid genome size of *Tubastraea* sp. to be about 1.1 Gb, corroborating the size estimated by flow cytometry, N50 of 227 kb in 7,996 contigs and completeness, as measured by BUSCO, of 91.6%. While the genome of *T. coccinea* was about 800 Mb, N50 of 66 kb in 17,214 contigs and 88.1% of completeness. The genome of *Tubastraea* sp. was also about 800 Mb and N50 of 82 Kb in 12,922 contigs and also 88,1% of completeness.

Our draft assemblies of *Tubastraea* spp. rank as the largest Cnidarian genomes published to date, twice to almost three times longer (800 Mb to 1 Gb vs 441±101 Mb) than published scleractinian genomes. The hybrid assembly strategy dramatically improved both contiguity and gene recovery of *T. tagusensis*, with the N50 increasing from about 6,000 bp to more than 220,000 bp, and BUSCO orthologous retrieval improving from about 50% (only short-reads) to more than 90%. We missed less than 10% of metazoan BUSCO genes in the genome and just 1% in the transcriptome. So this strategy was used to assemble the genome of *T. coccinea* and *Tubastraea* sp..

Intrinsic repetitive elements of the three morphotypes of *Tubastraea* sequenced in this work inferred using *ab-initio* and homology-based approaches with RepeatModeler and RepeatMasker, constituted about 50% of bases for *T. tagusensis*, and nearly 58% for *T. coccinea* and *Tubastraea* sp.. These percentages are higher than other species of Cnidaria group ^34,35^. Genome annotation using Breaker2 provided an estimate of around 121,000 genes for *T. tagusensis*, 51,000 for *T. coccinea* and 47,000 for *Tubastraea* sp.. The high gene content, almost 5-fold for *T. tagusensis* and 2-fold for *T. coccinea* and *Tubastraea* sp., the usual in Eukaryotes, could be attributed to reminiscent contamination ^40^, repeat masking, genome fragmentation ^41^, and warrants deeper scrutiny.

The functional annotation based on PANTHER protein families showed that the most representative pathways in *Tubastraea* sp. are: Wnt signaling; Integrin signalling; Gonadotropin-releasing hormone receptor pathway; Inflammation mediated by chemokine and cytokine signaling and Nicotinic acetylcholine receptor signaling. The Gene Ontology annotation showed a prevalence of protein families associated with biological regulation, cellular process localization and metabolic process.

To unveil the functional genomic idiosyncrasies of the *Tubastraea* genus in comparison to other Scleractinia and Cnidaria species we employed OrthoVenn 2. The gene models of *Nematostella vectensis* (OrthoVenn 2 built-in data), *Acropora digitifera* (GCF_000222465), *Orbicella faveolata* (GCF_002042975) and the predicted proteins for the three *Tubastraea* morphotypes were compared to identify orthologous genes and clusters. In a broader view, we identified 27,038 orthologous clusters which 26,770 are present in two or more species and 268 are single-copy gene clusters. The Cnidaria core-genome is composed of 17,263 proteins distributed on 1,759 gene sets and mainly related to basal activities such as biological regulation, metabolic process, and response to stimulus. The main molecular functions are associated with ion binding, peptidase, hydrolase and transferase activities, nucleic acid binding, and oxidoreductase activity. Regarding the core-genome of the Scleractinia order, we identified 6,075 proteins clustered on 554 orthologous sets. As for the Cnidaria core set, the Scleractinia clusters are mainly associated with basal functions.

Hard corals are marine pioneers and reef builders by excellence, then we searched for orthologous sets that might be involved in the aragonite skeleton structure and substrate adhesion. We identified seven private clusters comprising 133 proteins related to calcium metabolism and two clusters of fifteen orthologs linked to substrate adhesion. Interestingly, we could also identify in *Tubastraea* eleven clusters associated with cell-cell and cell-matrix adhesion that might be related to substrate affinity and runner-formation. Along with adhesion genes, proteins associated to the innate immunity might be of particular interest to the development of specific antifouling strategies for *Tubastraea* since incrustation is dependent on the establishment of a biofilm for larval settlement. Thus, we searched for enriched terms associated with innate immunity in *Tubastraea* and found orthologs sets associated with defense response to bacteria and TRIF-dependent toll-like receptor signaling pathways. Along with cell-matrix adhesion and immunity terms, the following functions are enriched in *Tubastraea*: sodium transmembrane transport, fatty acid biosynthetic process, regulation of kainate selective glutamate receptor activity, sensory receptors, signal transduction in response to DNA damage and RNA-mediated transposition. Most *Tubastraea* species-specific clusters are composed of uncharacterized proteins pointing to the pervasive knowledge gaps in corals and also highlighting their potential for bioprospection. In *T. tagusensis*, the term scavenger receptor activity is enriched and it is mainly associated with cell endocytosis or pattern recognition with a broad range of ligands which include Gram-positive and Gram-negative bacteria. We also identified gene-sets associated with spermatid development, tissue remodeling and regeneration, DNA repairing and response to radiation, heat response, hormone and protein precursors processing and cilium assembly. In *Tubastraea* sp., the serine-type endopeptidase activity term is enriched and it is related to collagen degradation, digestion, and antimicrobial peptides. Besides that, two more groups of orthologs are associated with innate immunity: one linked to tumor necrosis factor-mediated signaling pathway and the other to recognition of apoptotic cells. Interestingly, *T. coccinea* not only shares a higher amount of orthologs with other species than *Tubastraea* than *T. tagusensis* and *Tubastraea sp*. as it has more exclusive clusters than the other two Tubastraea analysed in this work. Gene Ontology enrichment test for *T. coccinea* pointed out the following terms: DNA integration, carbohydrate biosynthetic, acetylcholine catabolic process, and RNA-directed DNA polymerase activity. The exclusive gene content of *T. coccinea* is mainly related to response to stress, innate immunity, regeneration, regulation, and chromosome organization. In the next steps, we expect to better characterize the relationship of these findings with the invasive and opportunistic behaviour of *Tubastraea* species.

## Conclusion

We present one of the most comprehensive studies of a Scleractinia taxon, with the morphology, cytogenetics, transcriptome, mitochondrial and draft nuclear genome of three morphotypes of *Tubastraea*. They constitute the largest genomes of the Scleractinia order published to date. This study yielded findings which begin to fill some of the gaps in our understanding of the *Tubastraea* genus. The mitochondrial genome provides further evidence pertinent to the discussion of the species identification. Our morphological analysis and molecular phylogenetics of the mitochondrial genome and three gene markers of eighteen specimens show specific characters that support the occurrence of a new morphotype in the brazilian coast. We provided evidence of exclusive gene content related to Scleractinia and *Tubastraea* adhesion and innate immunity. These findings might unveil the biological basis of substrate affinity of hard corals, especially in pioneer species such as *Tubastraea* spp. and also provide support to the development of biotechnological antifouling strategies. Building upon the foundation of the work presented, in the next phases of our research we will improve the contiguity of the draft assembly by the use of molecular and computational scaffolding methods, elucidate the taxonomy within the *Tubastraea* genus by the use of integrative taxonomy, and generate better annotations to guide the development of biotechnological strategies to deter bioinvasion by this species of sun coral and to gain insights about resilience among native corals.

## Disclaimer

To enhance readability, we elected to organize this article in a narrative format. The content of the conventional scientific format of Introduction, Materials Methods, Results, and Discussion is present, just not with these subtitles. A detailed materials, methods and results section is presented after the references.

## Acknowledgments

Authors are indebted to Mrs Lisboa M for her support. .Dr. Viccini LF and Dr. Monteiro E for their support in flow cytometry. Dr. Zilberberg C for her insights in DNA extraction and Guimarães M and Wajsenzon IJR for their invaluable help in experiments performance.

## Material, Methods and Results

### 1. Colonies sampling and identification

To analyse the complete genome, we collected a morphotype of *T. tagusensis* and other of *T. coccinea* on vertical substrate at 7 meter depth by scuba diving off the rocky shores of Porcos Pequena Island in the municipality of Angra dos Reis, Rio de Janeiro state, southeast Brazil (23°3.229’S; 44°19.058’W) (Supplementary Figure 1). Further, it was also sampled a distinct morphotype of *T. coccinea* found in the Porcos Island on vertical substrate at 8 meter depth, Arraial do Cabo (22°57’56”S; 41°59’36”W) (Supplementary Figure 1). We carefully removed the healthy colony from the substrate and immediately transported it to the laboratory. After a period of acclimation, we transferred the colony to a 20-liter seawater-filled aquarium (pH = 8.2; T°C = 24) which was continuously aerated and subjected to a 12h:12h photoperiod.

### Morphological analysis

After DNA extraction, the coral skeleton was prepared for morphological analyses by placing the colony in a container filled with hypochlorite solution (NaOCl) for the removal of all soft tissues. The bleaching time was three days. Then, samples were rinsed with deionized water and put to dry for 24–48 h. Colony and coralites were photographed using a Nikon D750 digital camera and a Leica M205 FA magnifying glass, respectively.

Macroscopic characteristics of the colony were examined with a Leica M205 FA magnifying glass. Calipers were used to measure the diameter of the colony and calice, as the height of the polyps. Characteristics measured in our colony were compared with published descriptions of *Tubastraea* species ^42^; ^43^; ^21,43^.

### Mitochondrial DNA analysis

The whole mitochondrial genome of the three specimens of coral sequenced in this work were retrieved from their respective scaffold databases with BLASTn (Altschul et al., 1990) using the *T. tagusensis* (NC_030352.1) ^32^ sequence as query. A second assembly was carried out with NOVOplasty software ^44^ using RNA reads in order to confirm the indels, polymorphic sites and regions that were not recovered in the whole genome assembly. Gene prediction and annotation was done with GeSeq ^45^ setting it to circular and allowing transfer RNA prediction with tRNAscan v2.0.5 ^46^. The global alignment of the whole mitochondrial genomes and others 65 form Scleractinian species available in NCBI was done with MAFFT using the auto mode ^47^. The maximum likelihood (ML) tree was built with RAxML version 8.0.0 ^48^ using the Generalised time-reversible model (GTR) ^49^. Branch reliability was estimated performing 500 replicates of bootstrap.

As for mitochondrial marker sequencing, DNA was extracted using the methodology described by ^50^ or CTAB lysis buffer and organic extraction ^51^. A region from the genes ATP8 and cytochrome oxidase I (COI) was amplified using the primers CS-16 described by ^52,53^. Another region between the 3’-end of the COI and 5’-end of the 16Sr ribosomal RNA (*rnl*) (IGR) was covered by the primer CS-18 ^52,54^ and a primer pair covering the cytochrome oxidase b (CytB) polymorphisms were used (Supplementary Table 1). The PCR reactions were carried out in a 25 µL reaction final volume containing 1X DreamTaq master mix (Thermo Scientific), 0.4 µM forward primer, and 0.4 µM reverse primer and 100 ng of DNA (CS-18 and CytB primers) or 500 to 750 ng (CS-16 primer), following cycling conditions: Initial denaturation at 95ºC for 3 minutes, followed by 35 cycles of denaturation at 95ºC for 30 seconds, primer annealing at 48ºC (CS-16) or 55ºC (CYTB and CS-18) for 30 seconds, amplification at 72ºC for 1:15 minutes, and final extension at 72ºC for 10 minutes. After visualization in agarose gel, the three markers amplified successfully in eighteen specimens, whose amplicons were sent to Macrogen Inc. (Seoul, South Korea), Helixxa (São Paulo, Brasil) or Senai Cetiqt (Rio de Janeiro Brasil), purified and sequenced by Sanger sequencing technique with the forward and reverse primers.

All forward and reverse chromatograms were manually inspected using SnapGene Viewer (SnapGene viewer). Only peaks with quality higher than 20 with non-overlapping signals were considered. All sequences of each marker were independently aligned with MAFFT using the G-INS-i strategy and concatenated in a single alignment with SeaView ^55^. The neighbor joining tree was built using Kimura’s 2-parameters ^56^ with 1000 replicates of bootstrap. The ML tree was built with RAxML version 8.0.0 using GTR model + GAMMA model of rate heterogeneity. To estimate the branch reliability 500 replicates of bootstrap were performed. Tree visualization and needed editions of the trees were done with Itol ^57^.

**Supplementary Figure 1:**
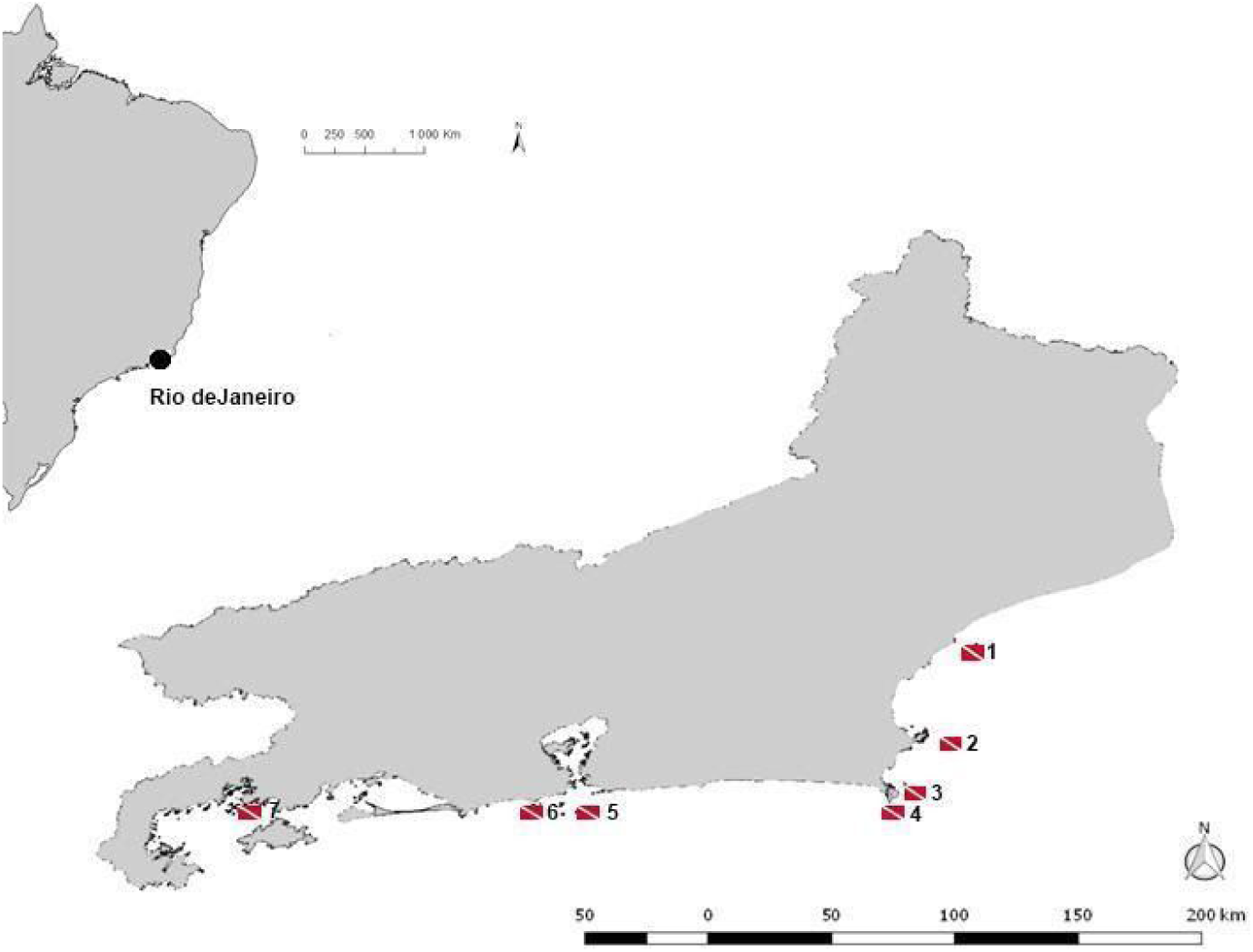
Map of sampled sites at Rio de Janeiro’s coast: (1) Francês Island – Macaé; (2) Âncora Island - Armação de Búzios (22°46’08”S; 41°47’12”); (3) Papagaio Island – Cabo Frio (22°54’00”S; 41°59’00”W); (4) Porcos Island - Arraial do Cabo (22°57’56”S; 41°59’36”W); (5) Filhote da Cagarra Island - Rio de Janeiro (23°01’52”S / 43°11’35”W); (6) Ilhas Tijucas - Rio de Janeiro (23°01’57”S / 43°18’05”W); (7) Porcos Pequena Island - Angra dos Reis (23°3.229`S / 44°19.058`W).

**Supplementary Figure 2.**
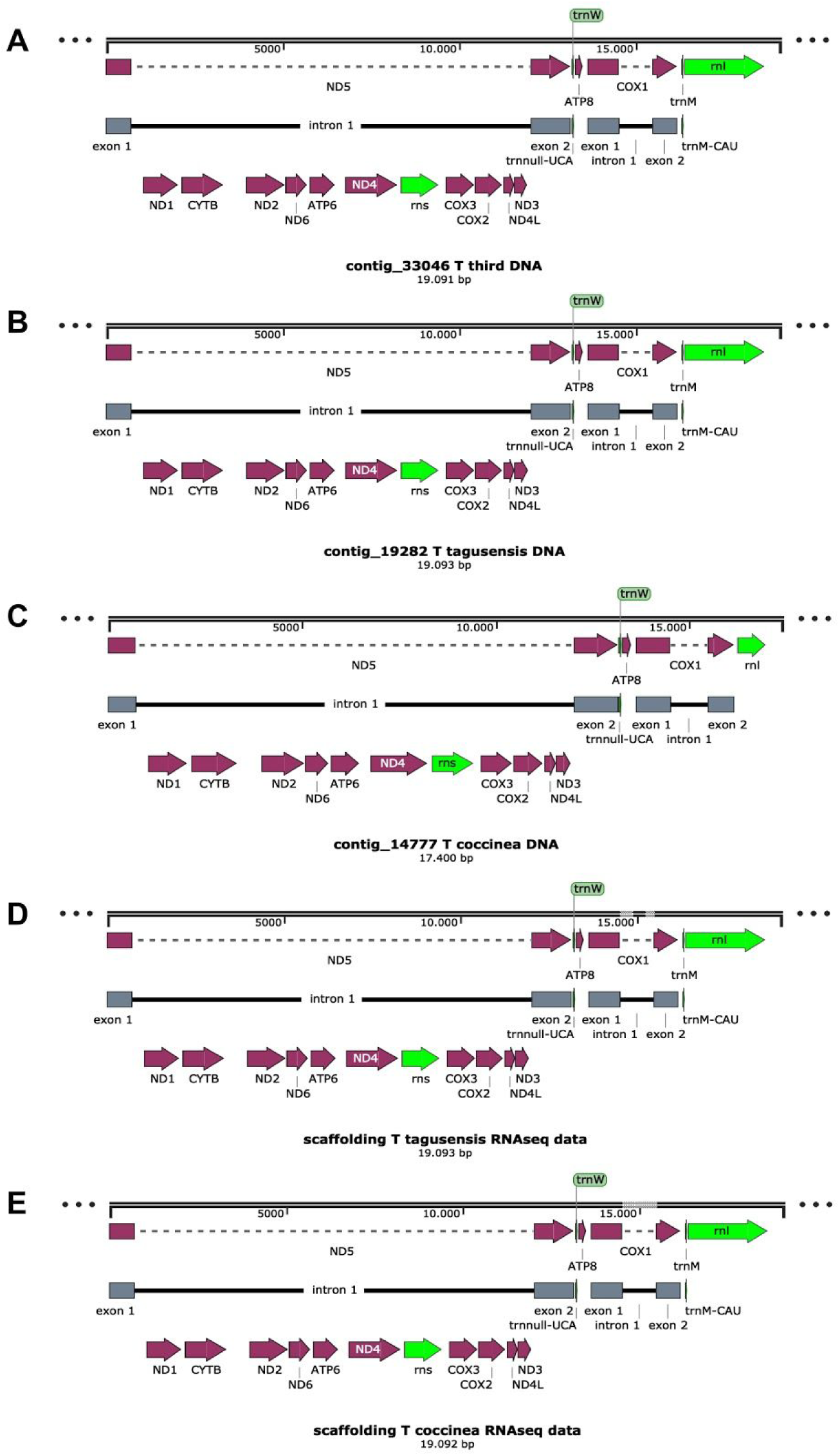
Mitochondrial genome annotation. Genes that encode for proteins are coloured in purple and ribosomal subunits genes in green. tRNAs tmW and tmM are shown before ATP8 and rnl genes, respectively. Exons of the ND5 and COX1 genes are coloured in grey. Introns are shown as solid black lines. A - *Tubastraea sp.* (DNA assembly; contig 33046). B - *T. tagusensis* (DNA assembly; contig 19282). C - *T. coccinea* (DNA assembly; contig 14777). D - *T. tagusensis* (RNA assembly). E - *T. coccinea* (RNA assembly).

**Supplementary Table 1:**
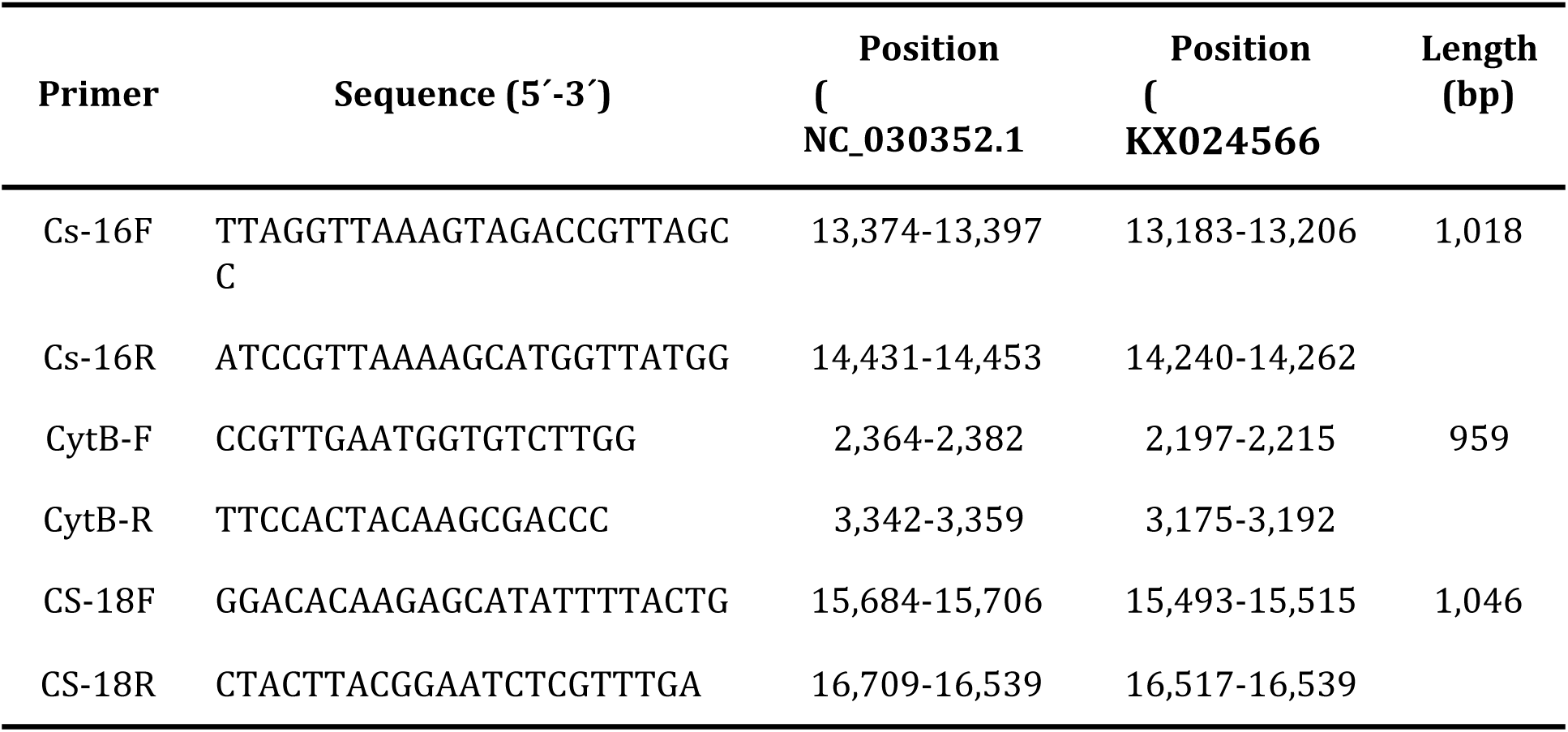
Sequence of the primers used to amplify mtDNA fragments from *Tubastraea* spp.

### 2. Flow cytometry

The genome sizes of *Tubastraea* spp. were estimated by flow cytometry. For the nuclei suspension preparation, a polyp was sampled and a 1 mm piece was minced in a buffer containing 0.2 M Tris-HCl, 4 mM MgCl_2_.6H_2_O, 2 mM EDTA Na_2_·2H_2_O, 86 mM NaCl, 10 mM sodium metabisulfite, 1% PVP-10, 1% (v/v) Triton X-100, pH 7.5. *Pisum sativum* (pea) was chopped in the same buffer and used as an internal standard for genome size estimation. The nuclear suspension was stained with propidium iodide and at least 5,000 events were analyzed using CytoFLEX (Beckman Coulter Life Sciences). Histograms were analyzed using CytExpert 2.0 software (Beckman Coulter Life Sciences). The 2C DNA content was calculated as the sample peak mean, divided by the *P. sativum* peak mean and multiplied by the amount of *P. sativum* DNA (9.09 pg, ^58^). The procedure was performed in experimental quadruplicate for T. tagusensis and experimental triplicate to *T. coccinea* and *Tubastraea* sp. The estimated 2C DNA content and genome sizes are shown in Supplementary Table 2 and representatives histograms are shown in Supplementary Figure 3.

**Supplementary Figure 3:**
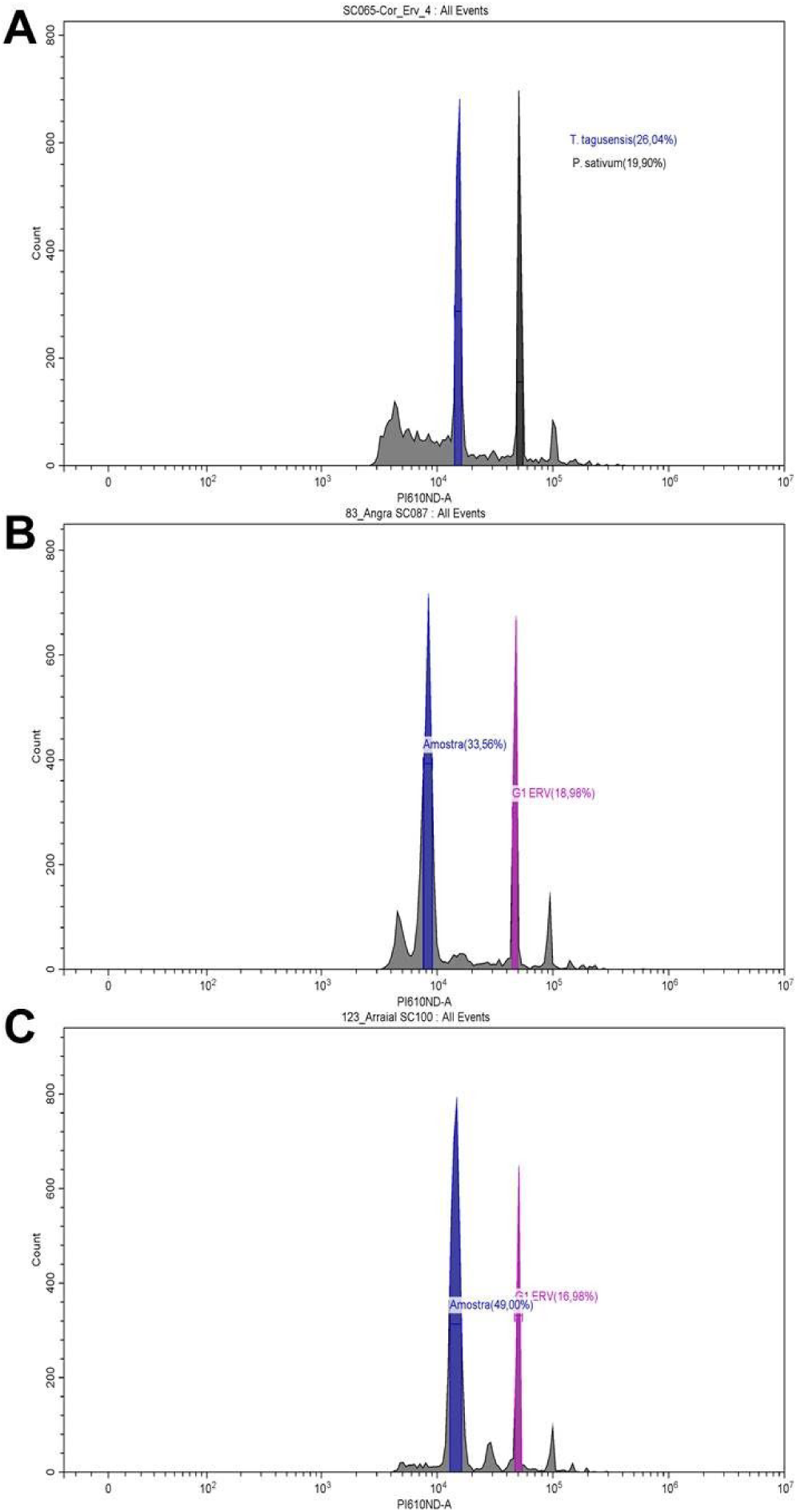
Representative histograms of DNA staining in nuclei with propidium iodide. In black or pink nuclear DNA fluorescence of the standard *P. sativum* (pea) and in blue fluorescence of *Tubastraea* spp. The 2C nuclear DNA content of **(A)** *T. tagusensis* was 2.61 pg (± 0.78), equivalent to 2,555 Mb (±77) and the genome size was estimated at 1,277 Mb (± 82). The nuclear DNA content of **(B)** *T. coccinea* was 2.35 pg (± 0.03), equivalent to 2,294 Mb (±22), and the genome size was estimated at approximately 1,147 Mb (± 15). And **(C)** the nuclear DNA content of *Tubastraea* sp. was 2.75 pg (±0.16), equivalent to 2,690 Mb (± 163), and the genome size was estimated in approximately 1,345 Mb (± 82).

**Supplementary Table 2:** *Tubastraea* spp. DNA content estimated by flow cytometry.

|  | *T. tagusensis* | *T. coccinea* | *Tubastraea* sp. |
| --- | --- | --- | --- |
| 2C DNA content (pg) | 2.61 ($\pm 0.78$) | 2.35 ($\pm 0.03$) | 2.75 ($\pm 0.17$) |
| 2C DNA content (Mb) | 2,555 ($\pm 77$) | 2,294 ($\pm 22$) | 2,690 ($\pm 163$) |
| Genome size (Mb) | 1,277 ($\pm 82$) | 1,147 ($\pm 15$) | 1,345 ($\pm 82$) |

### 3. Karyotype determination

Planulae from *Tubastraea* spp. were collected for karyotype determination. Planulae were exposed to colchicine (0.1%) for 24h, then exposed for 90 minutes to an osmotic shock with distilled water to rupture the membranes, and then fixed with Carnoy’s solution (3:1; Ethanol: acetic acid). The fixed planulae were then minced in acetic acid (60%), placed onto a heated glass slide and air-dried. Nucleic acids on the slide were stained with DAPI (4′,6-diamidino-2-phenylindole) for visualization under fluorescence microscopy. Chromosomes were measured using Adobe Photoshop (Adobe Systems) and was possible to observe 46 chromosomes in *T. tagusensis* 44 chromosomes in *T. coccinea* and 42 chromosomes in *Tubastraea* sp. (Figure 2, Supplementary Figure 4).

**Supplementary Figure 4:**
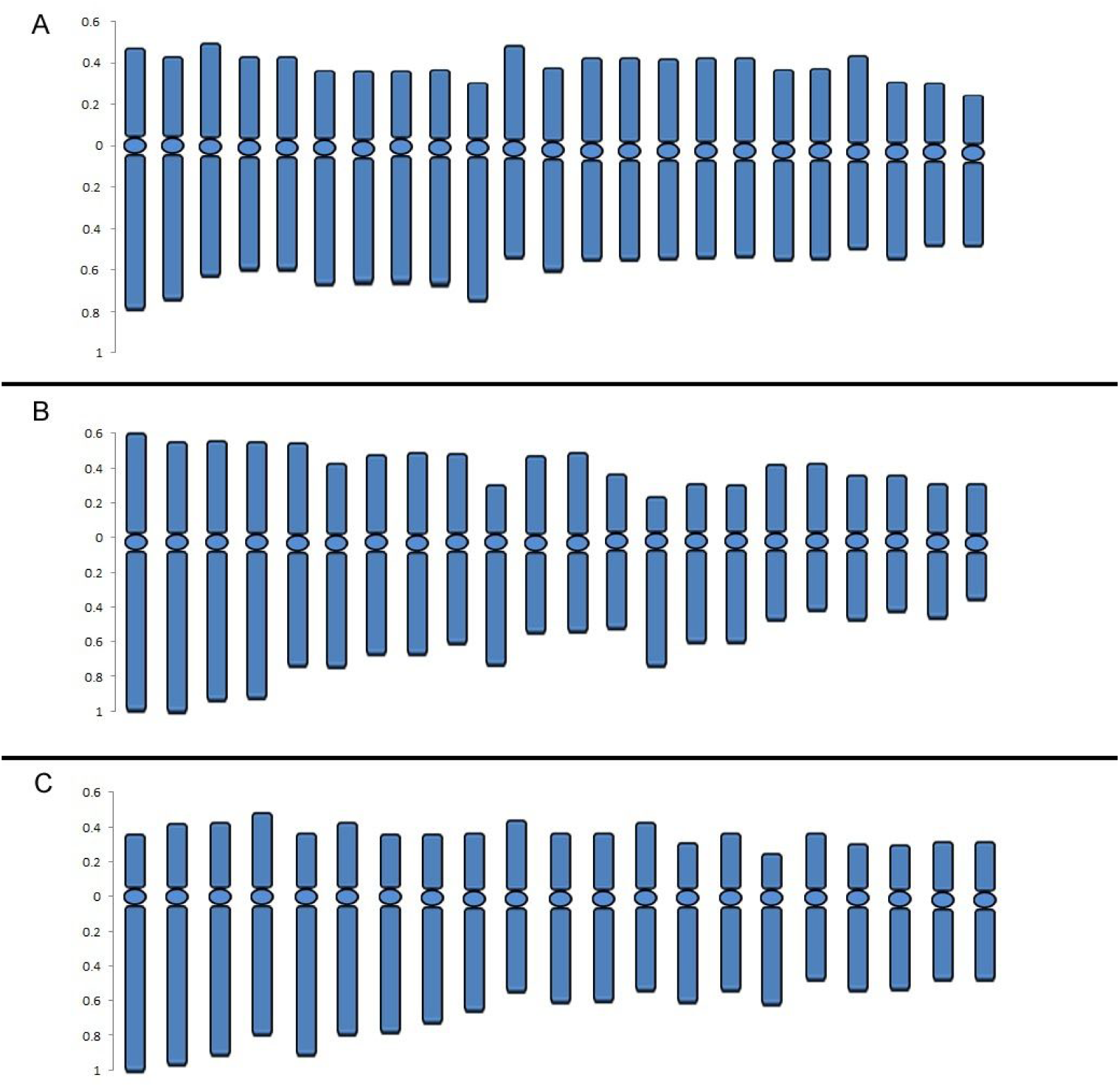
Ideogram of (A) *T. tagusensis* (2n= 46), (B) *T. coccinea* (2n= 44), and (C) *Tubastraea* sp. (2n= 42).

### 4. DNA and RNA extraction

Soft tissue from the coral specimen was collected, rinsed with distilled water and DNA extraction was performed. The DNA extraction from *T. tagusensis* destined for Illumina sequencing was performed using the *DNaesy* Blood & Tissue kit (Qiagen). The DNA extraction from *T. tagusensis* destined for PacBio sequencing and *T. coccinea* and *Tubastraea* sp. destined for both illumina and PacBio sequencing was performed according to ^50^, optimized with several modifications.

Briefly, soft tissue from sun coral specimens retrieved from aquaria were rinsed with distilled water and immersed in 1.0 ml of CTAB buffer [2 % (m/v) CTAB (Sigma-Aldrich), 1.4 M NaCl, 20 mM EDTA, 100 mM Tris–HCl (pH8,0)], with 10 µg of proteinase K (Invitrogen) and 2% of 2-mercaptoethanol (Sigma-Aldrich), freshly added, per 100 mg of tissue. Then, the tissue was kept in a lysis buffer for 4 days, with occasional inversion to promote tissue lysis.

Tubes with tissue and buffer were then exposed to freezing with liquid nitrogen for 30 seconds and then thawed and heated to 65ºC in a heat block for approximately 3 minutes. Three freeze-thaw cycles were performed. Then to remove protein and lipids one wash was performed with Phenol:Chloroform:Isoamyl Alcohol (25:24:1) (Sigma-Aldrich), and two washes with Chloroform:Isoamyl Alcohol (24:1) (Sigma-Aldrich). The supernatant was transferred to a new tube containing 1 ml of C4 solution from a Power Soil DNA Isolation kit (MO BIO Laboratories), homogenized by inversion and loaded in a spin column from the *DNeasy* Blood & Tissue kit (Qiagen, Germany). A final wash was performed with 500 µl of C5 solution. DNA elution was done with 150 µl of Tris-HCl buffer (10 mM, pH 8.5) in three sequencing centrifugation steps. RNA extraction from *Tubastraea* sp. from Angra dos Reis was done using the Trizol protocol, according to the manufacturer’s instructions with a few modifications. A polyp of each species was homogenized with a rotor-stator in 1 ml of TRIzol (Thermo Fisher) or TRI-Reagent (Merck) with 6 µl of HCl (6M) ^59^. The homogenate was centrifuged to remove debris and then we followed the Trizol or TRI-Reagent manufacturer’s protocol.

Both DNA and RNA purity was assessed using a NanoDrop spectrophotometer (Thermo Fisher Scientific), the amount determined using a Qubit Fluorometer (Thermo Fisher Scientific). DNA and RNA integrity was evaluated by 1.0% agarose gel electrophoresis and the respective ScreenTape Assay using a 4200 Tapestation System (Agilent).

DNA extraction of *T. tagusensis* using the *DNeasy* Blood & Tissue kit yielded a DNA of about 10 kb in size with a DNA Integrity Number (DIN) of 6.4 (Supplementary Figure 5) and free of proteins and was used to construct the Illumina library. DNA extraction using the CTAB buffer protocol yielded a DNA of about 23 kb in size and DIN = 8.0 (Supplementary Figure 6) and also free of proteins and other contaminants. It was used to construct the SMRTbell library. The DNA extraction from *T. coccinea* yielded a DNA > 60 kb in size and DIN = 8.0 (Supplementary Figure 7). And the DNA extraction from *Tubastraea* sp. yielded a DNA around 14 kb in size (Supplementary Figure 8).

**Supplementary Figure 5:**
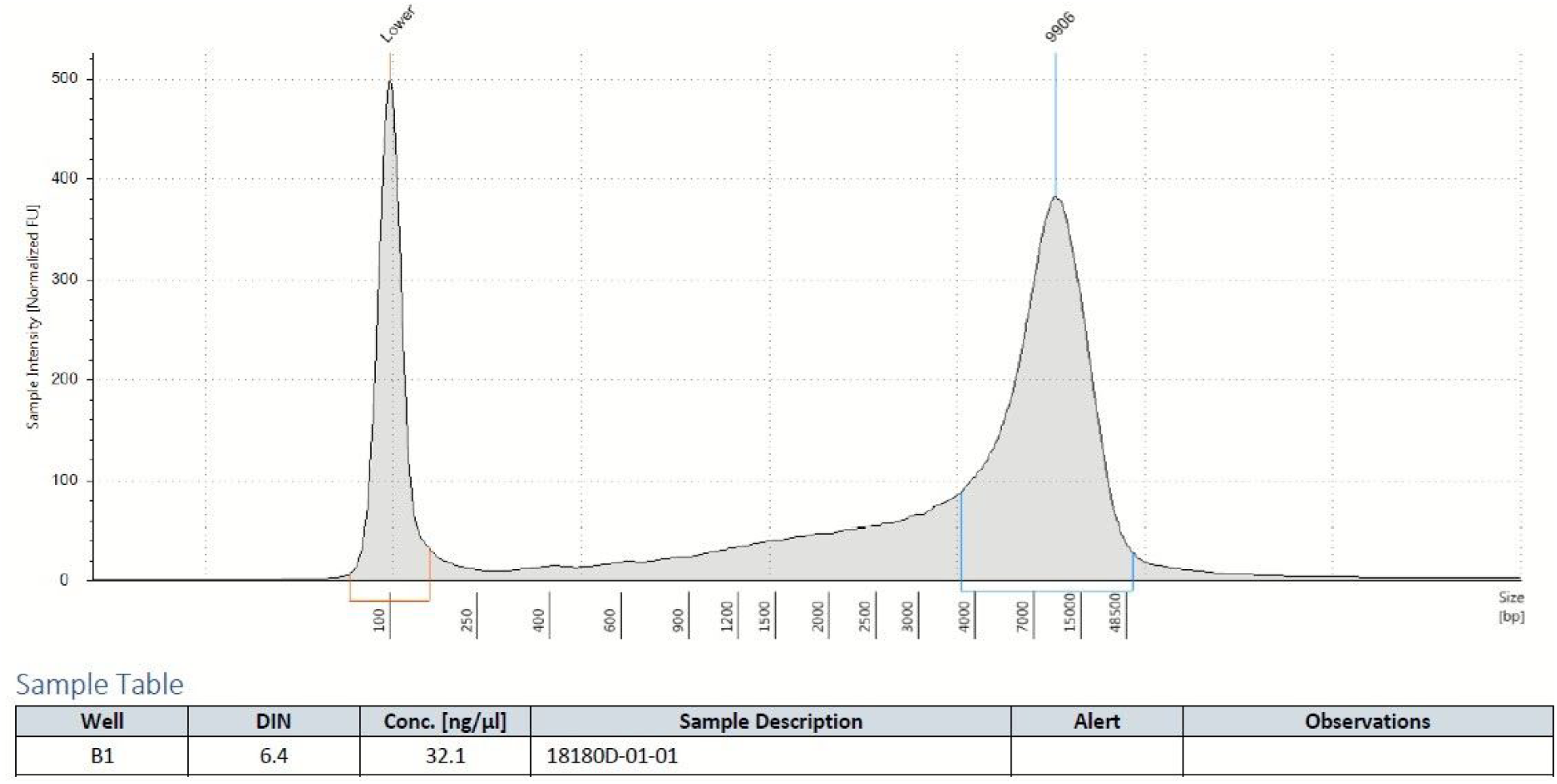
TapeStation evaluation of the DNA from *T. tagusensis* extracted with DN*easy* Blood & Tissue kit.

**Supplementary Figure 6:**
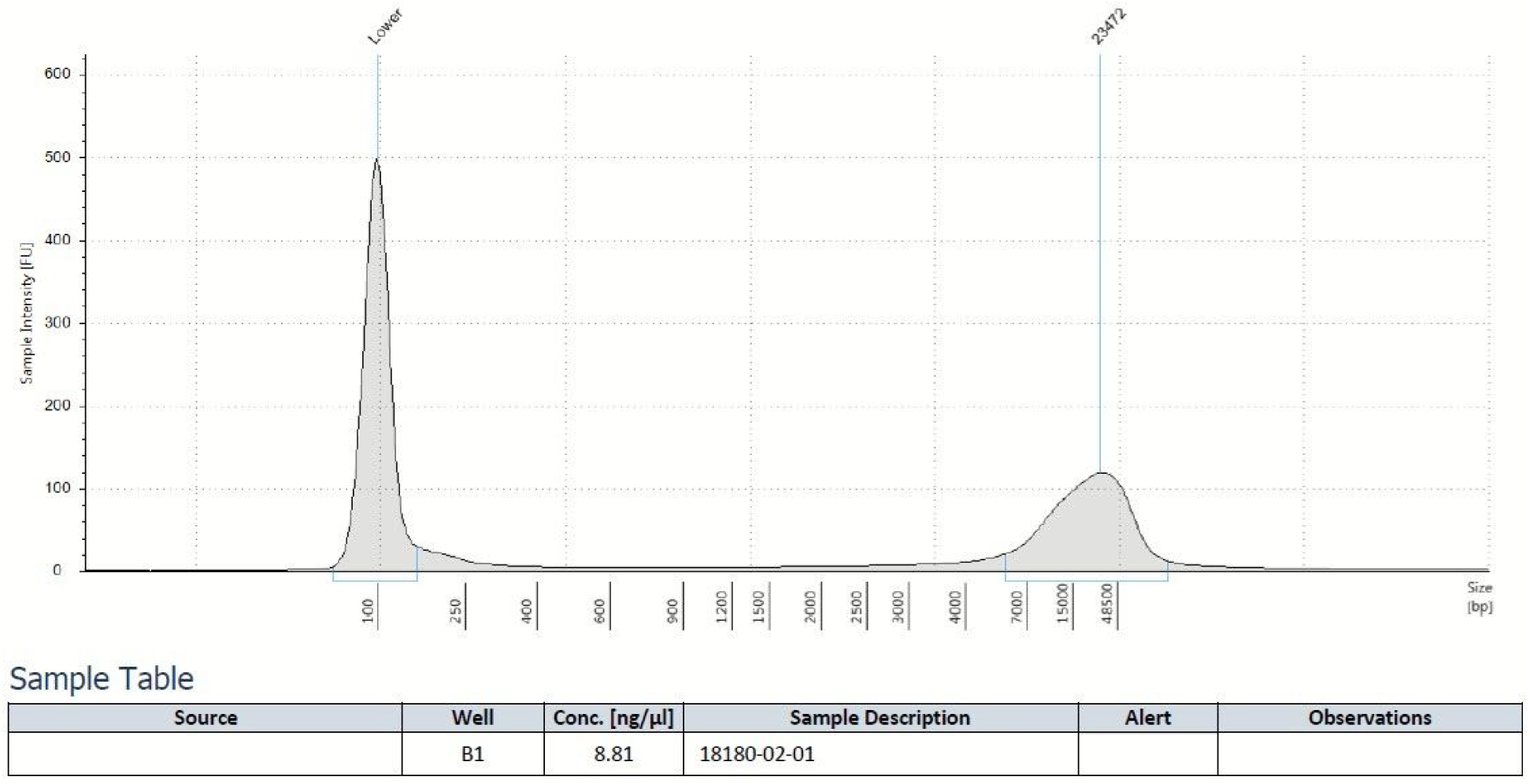
TapeStation evaluation of the DNA from *T. tagusensis* extracted with CTAB buffer combined with MoBio PowerSoil kit.

**Supplementary Figure 7:**
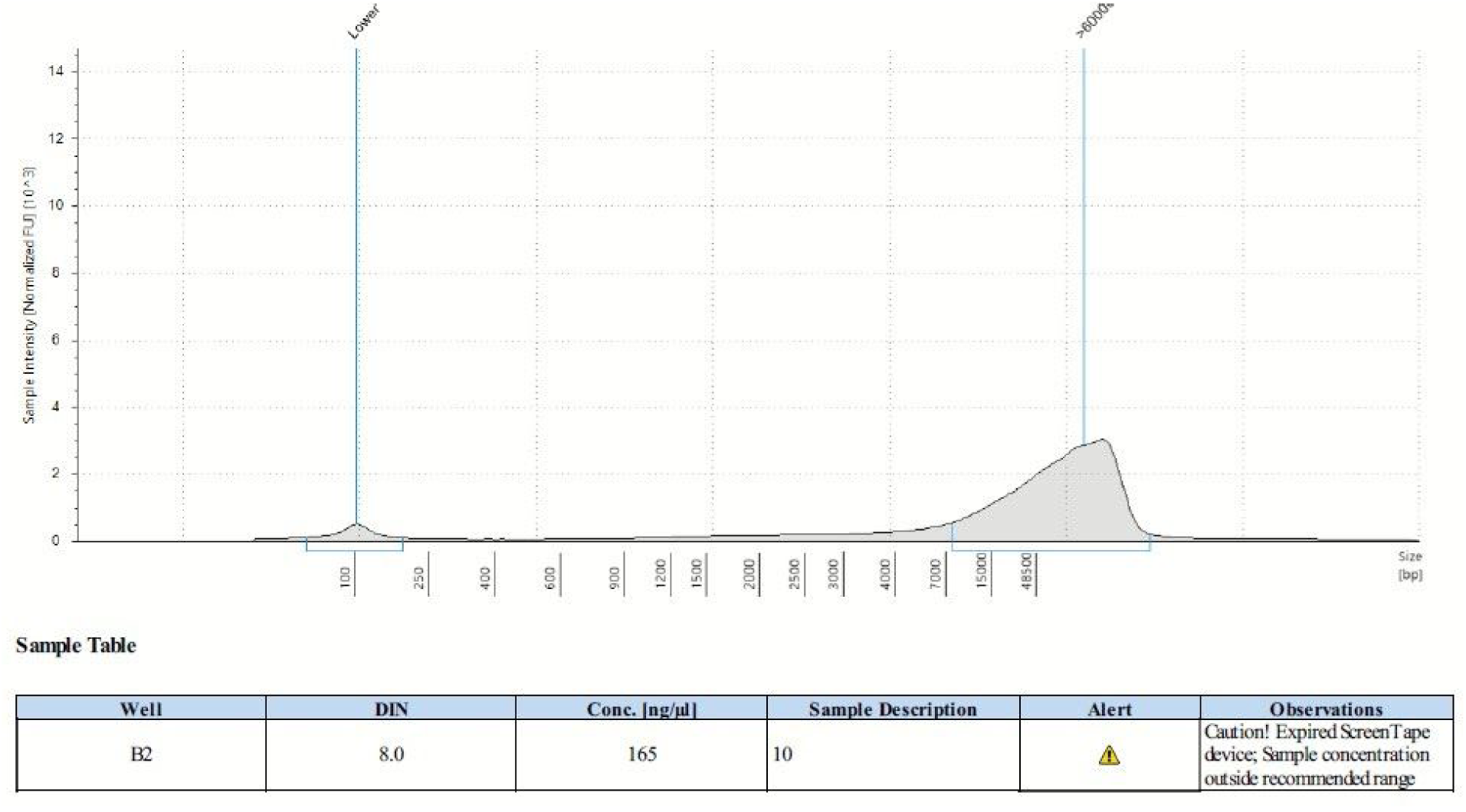
TapeStation evaluation of the DNA from *T. coccinea* extracted with CTAB buffer combined with MoBio PowerSoil kit.

**Supplementary Figure 8:**
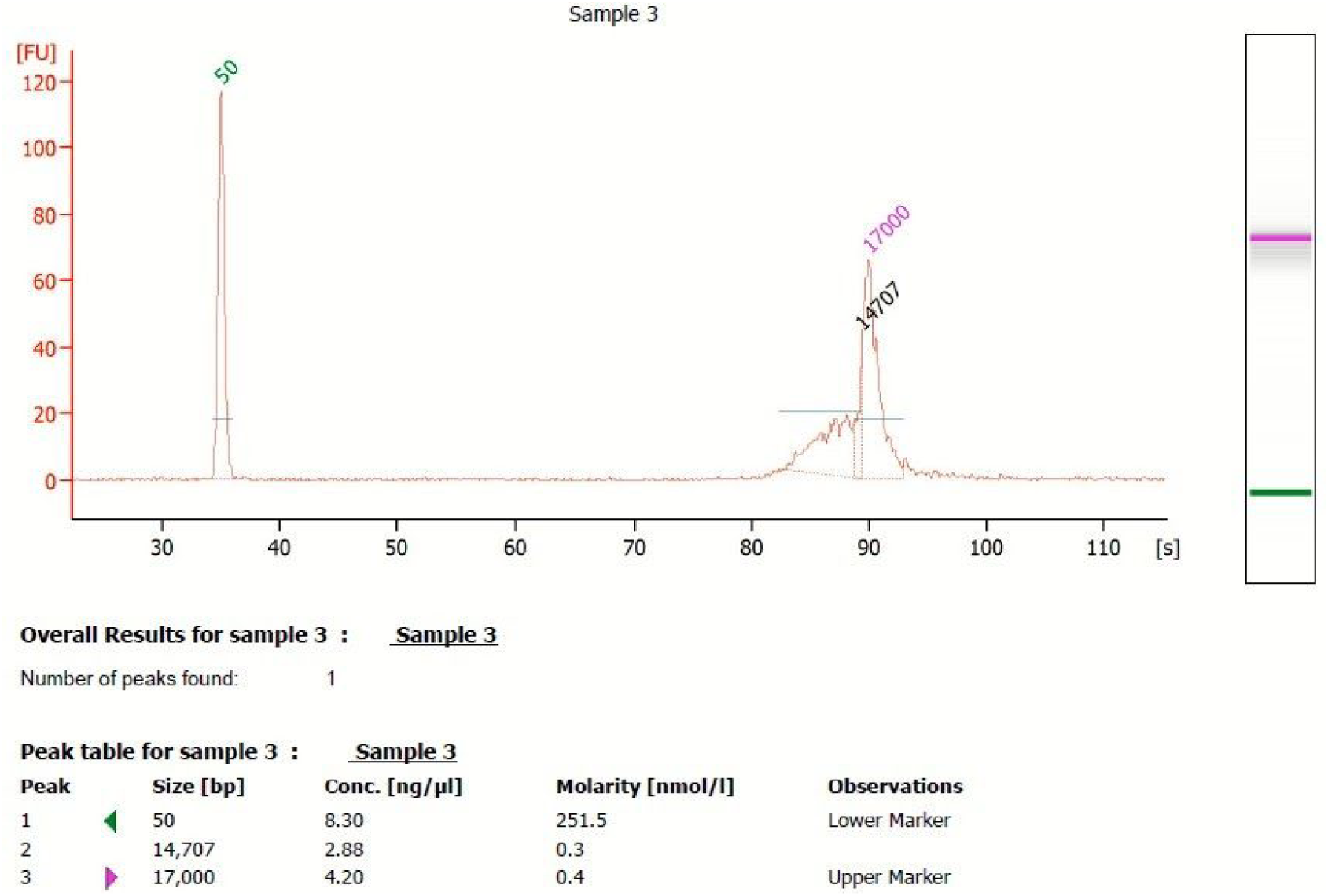
TapeStation evaluation of the DNA from *Tubastraea* sp. extracted with CTAB buffer combined with MoBio PowerSoil kit.

RNA extraction yielded RNA with quality enough for library construction. The RNA integrity number (RIN) of *T. tagusensis* was 6.6, of *T. coccinea* was 7.1, and of *Tubastraea* sp. 8.2 (Supplementary Figure 9). NEBNext mRNA libraries were built using magnetic isolation and 30 million 150 bp paired-end reads were sequenced using an Illumina HiSeq X.

**Supplementary figure 9:**
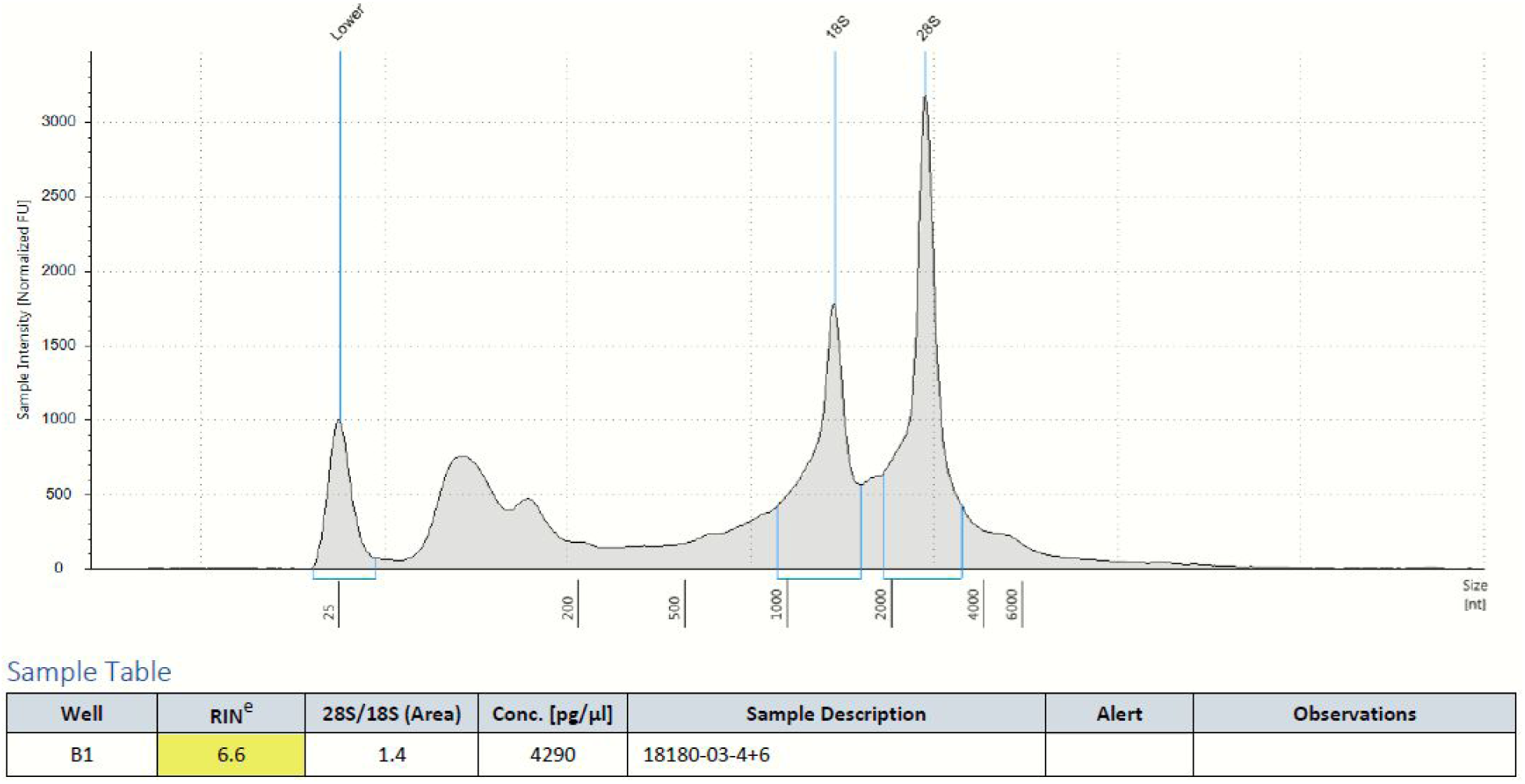
Electropherogram and RNA Integrity Number (RIN) of *T. tagusensis* RNA.

**Supplementary Figure 10:**
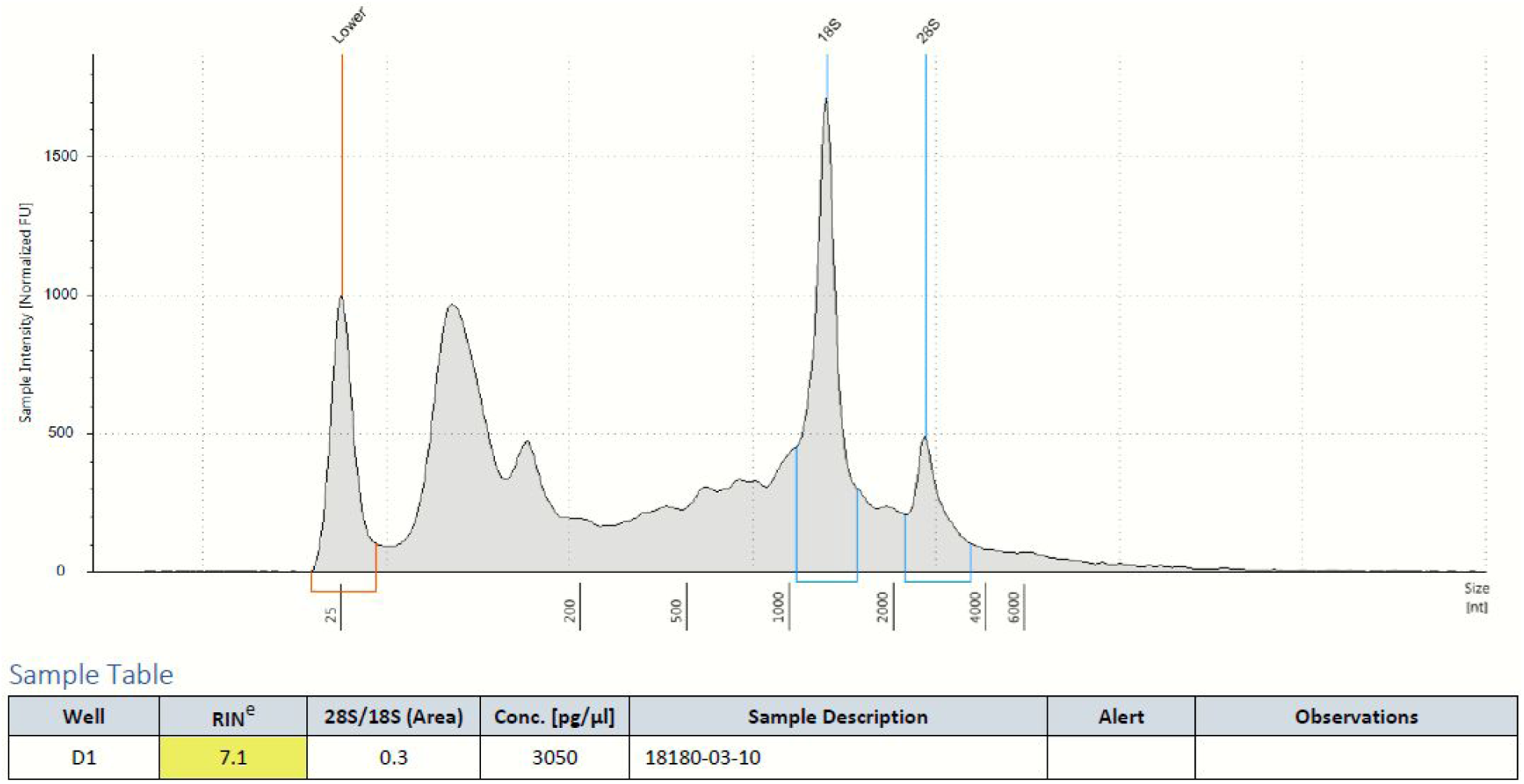
Electropherogram and RNA Integrity Number (RIN) of *T. coccinea* RNA.

**Supplementary Figure 11:**
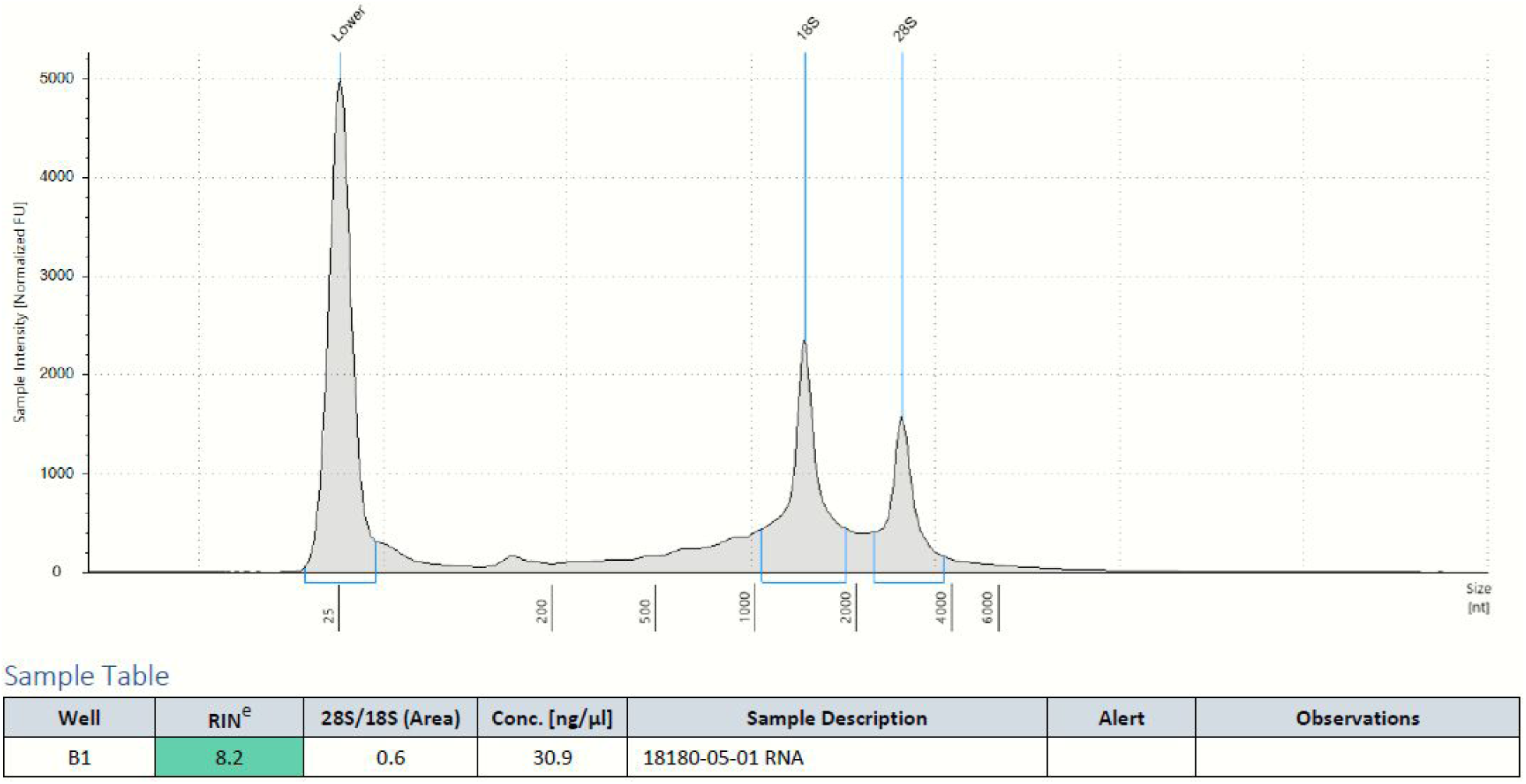
Electropherogram and RNA Integrity Number (RIN) of *Tubastraea* sp. RNA.

### 5. RNA sequencing and annotation

#### Library construction and sequencing

Library construction of *T. tagusensis, T. coccinea* and *Tubastraea* sp. were performed following the manufacturer’s recommendations for the NEBNext Ultra II RNA Library Prep Kit for Illumina with the NEBNext Poly(A) mRNA Magnetic Isolation Module (New England Biolabs, Ipswich, MA). Samples were pooled and sequenced on a HiSeq X sequencer at a 150 bp read length in paired-end mode, with an output of 30 million reads per sample.

#### Transcriptome Quality control

The stranded paired-end RNA sequencing raw data was first evaluated using fastQC v.0.11.8 and KAT v.2.4.1. After the visual inspection of the graphs and statistics generated, bbDUK v.38.42 was used to perform the quality trimming and filtering of reads. Illumina PE data was modulo trimmed (ftm = 5) due to the lower quality of the last base in the 151 bp reads. In the second round, reads were quality trimmed using the following parameters: i) minlength=100 (minimum length); ii) ktrim=r (trim reads matching reference k-mers); iii) k=23 (k-mer size used to find contaminants); iv) hdist=1 (Hamming distance for reference k-mers); v) tbo=t (trim adapters based on reads overlap); vi) tpe=t (trimming on both reads); vii) qtrim=rl (trim both reads ends based on quality score); viii) trimq=20 (quality trimming score); and ix) minavgquality=20 (minimum average read quality). Read contaminants were searched against phix and human sequences.

#### Transcriptome Assembly

Trinity v.2.8.5 was used in the *de novo* transcriptome assembly of *Tubastraea* spp. with the following parameters: --seqType fq --max_memory 50G --CPU 16. The quality of transcriptome assembly was evaluated by different metrics such as: i) transcriptome statistics; ii) reads mappability in the transcriptome and the genome; iii) proportion of full-length reconstructed protein-coding genes from predicted transcripts; iv) proportion of recovered conserved orthologous genes.

Trinity v.2.8.5 stats script was used to calculate the summary statistics of transcriptomes assemblies. Bowtie2 v.2.3.5 were used to map the reads to the transcriptomes with the following parameters (-p 16 -q --no-unal -k 20 -x) and 75% of them of each morphotype aligned to the genome using STAR v.2.7.2a with default parameters.

To estimate the number of protein-coding genes represented by putative transcripts we carried out the following steps: i) “blasted” (blastx v.2.9.0+) the putative transcripts against the SwissProt database release 2019_07 (-evalue 1e-20 -num_threads 16 -max_target_seqs 1 -outfmt 6); ii) estimated the proportion of protein sequence targets aligned to the best match transcript (analyze_blastPlus_topHit_coverage.pl); iii) grouped multiple high scoring segment pairs (HSPs) hits to estimate sequence coverage based on multiple alignments (blast_outfmt6_group_segments.pl); iv) computed the percent coverage by length distribution. We also computed the number of predicted orthologous genes recovered from transcriptome assembly using BUSCO v3.0.2 (Benchmarking Universal Single-Copy Orthologs) with default flags (-m transcriptome -c 32 -sp fly; metazoa database). The summary statistics of the transcriptomes are shown in Supplementary Table 3.

#### Transcriptome Annotation

TransDecoder.LongOrfs v5.5.0 with default parameters was used to identify the most likely candidate regions in assembled transcripts. The predicted open reading frames (ORFs) were then searched using blastp v2.9.0+ (-outfmt 6 -evalue 1e-5 -num_threads 20 -outfmt 6 -evalue 1e-5 -num_threads 20 -max_target_seqs 1) against the SwissProt database release 2019_07 and using hmmscan v3.2.1 against the Pfam database v32.0 to create a homology retention filter for TransDecoder.Predict.

**Supplementary Table 3:** Summary statistics of transcriptome assembly.

| **Transcriptome**<br>**Assembly**<br>**Information** | ***T. tagusensis*** | ***T. coccinea*** | ***Tubastraea* sp.** |
| --- | --- | --- | --- |
| Alignment rate | 94.81% | 88.94% | 93.08% |
| Trinity genes | 208,419 | 259,653 | 117,131 |
| Trinity transcripts | 357,670 | 392,794 | 196,392 |
| Transcript contig N50 | 1,407 | 850 | 1,270 |
| Transcript average<br>length | 803.13 | 593.79 | 787.04 |
| BUSCO Complete | 98.10% | 80.2% | 91.8% |
| BUSCO Fragmented | 1.30% | 4.3% | 6.2% |
| BUSCO Missing | 0.60% | 15.5% | 2% |
| Full length transcripts<br>(Uniprot > 80%) | 5,638 (39%) | 4,171 (28%) | 4,196 (35%) |

### 6. DNA sequencing and annotation

#### Illumina Library construction and sequencing

The DNA library from *Tubastraea tagusensis.* was prepared using the Kapa Hyper DNA Library Preparation Kit, with a final library size of 494 pb (Supplementary Figure 12). The 150 bp paired-end library was sequenced on a HiSeq X (Illumina, Inc., San Diego, US-CA).

**Supplementary Figure 12:**
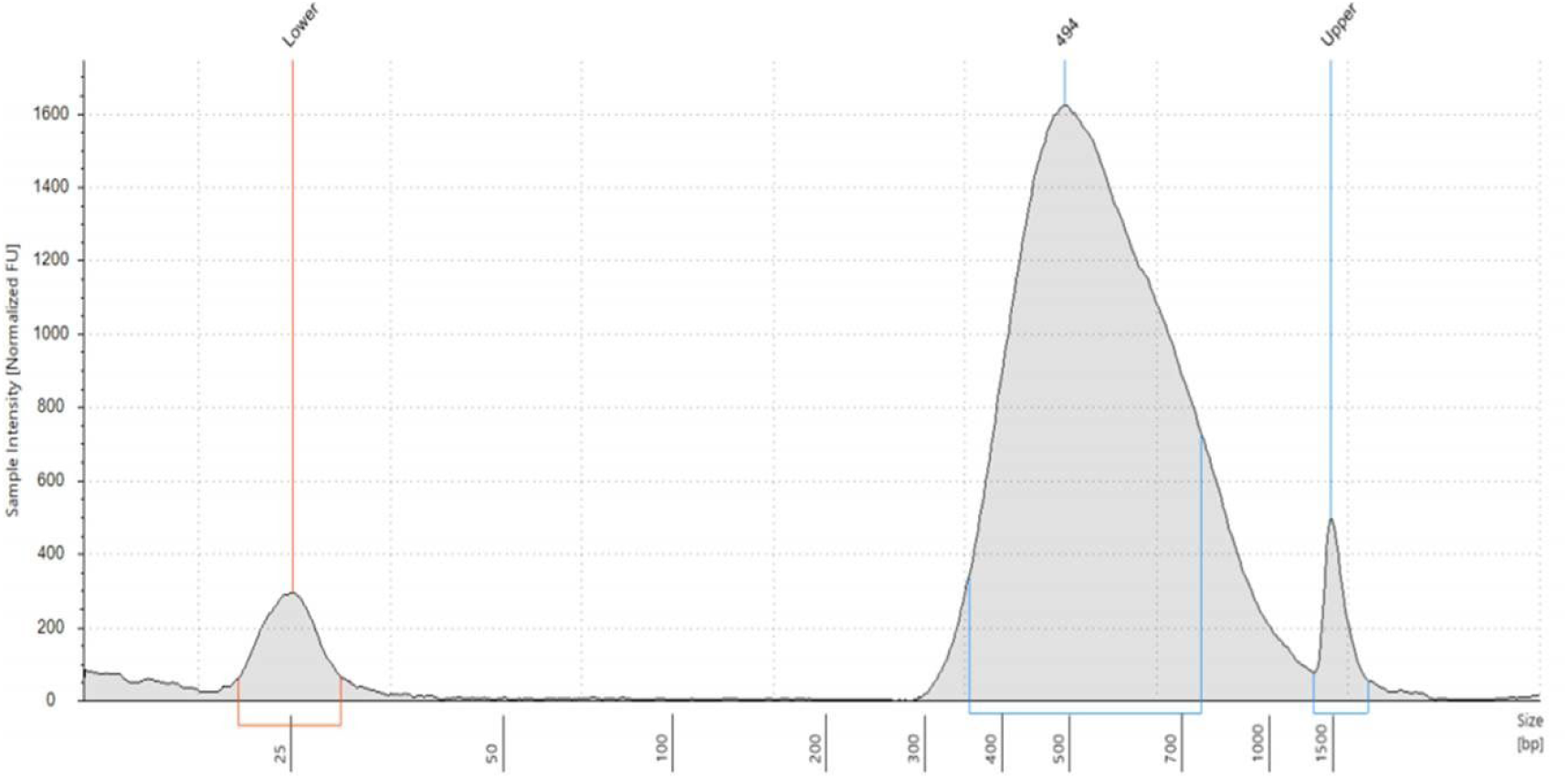
*Tubastraea tagusensis* library size evaluation. Electropherogram traces of the constructed Illumina library.

A total of 500 ng of DNA from *T. coccinea* and *Tubastraea* sp. was used to construct the library for sequencing using Illumina technology. The libraries were constructed using the Nextera DNA Flex Library Prep Kit. The sequencing platform used was the Illumina NextSeq 550, with the NextSeq 550Dx High Output Reagent Kit v2 (300 cycles). The average size of the libraries of *T. coccinea* and *Tubastraea* sp. were 557 bp (Supplementary Figure 13) and 496 bp, respectively (Supplementary Figure 14).

**Supplementary Figure 13:**
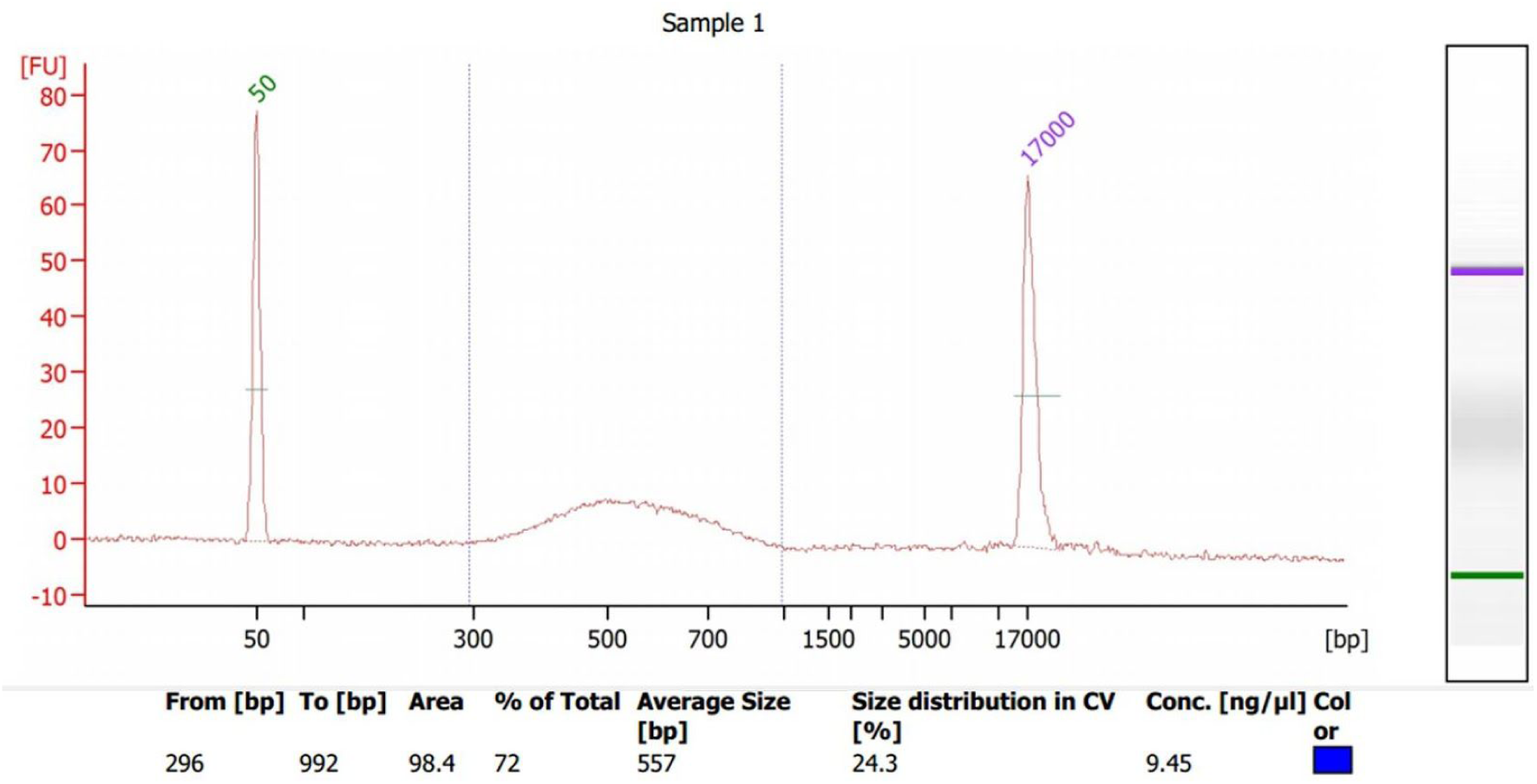
*Tubastraea coccinea* library size evaluation. Electropherogram traces of the constructed Illumina library.

**Supplementary Figure 14:**
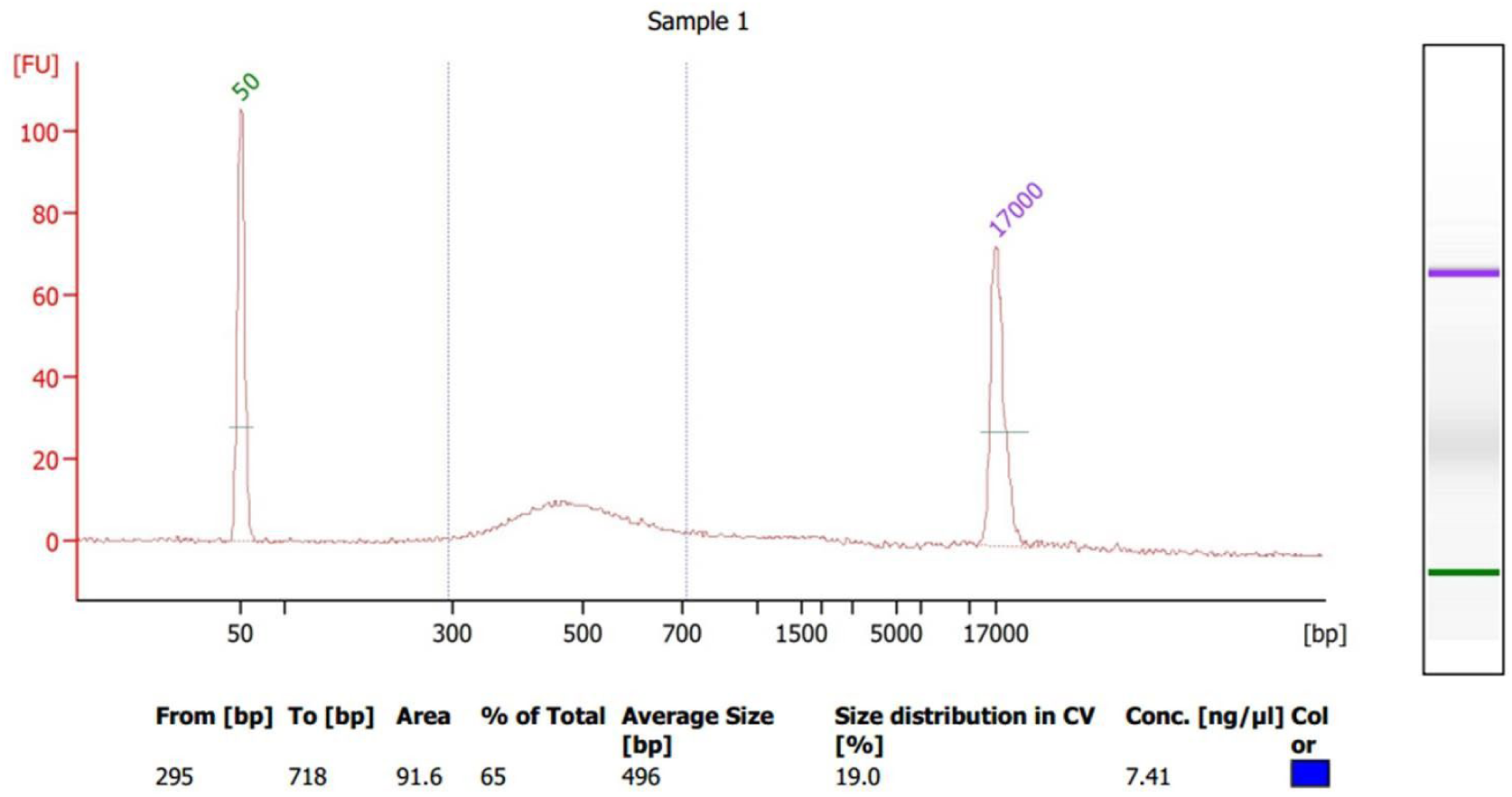
*Tubastraea* sp. library size evaluation. Electropherogram traces of the constructed Illumina library.

### PacBio library construction and sequencing

#### Tubastraea tagusensis

As for the SMART bell library preparation, 5 µg of total DNA of *T. tagusensis* were processed without further shearing the DNA as it was already measured to be around 20Kb. After the steps to repair the DNA repair and to ligate adapters and DNA that didn’t have SMRTbell primers attached were removed, then 3 µg were loaded onto Bluepippin equipment, using S1 markers (Sage Science) as standard. Everything at the size range >15kb were collected to make sure there was enough material recovered to load at the Sequel. The average size of the library was 15kb (data not shown).

#### Tubastraea coccinea

As for the SMART bell library preparation, 15 µg of total DNA of *T. coccinea* were processed and sheared with Covaris g-TUBE. After the steps outlined in the attached protocol to repair the DNA and ligate adapters and then remove DNA that didn’t have SMRTbell primers attached. Then DNA was purified using 0.45xAMPure purification and fragments <1.5 kb were removed using 0.40xAMPure purification. Then, 4 pM was used to load the SMRT cell. The average size of the library was 12 kb (Supplementary Figure 15).

**Supplementary Figure 15:**
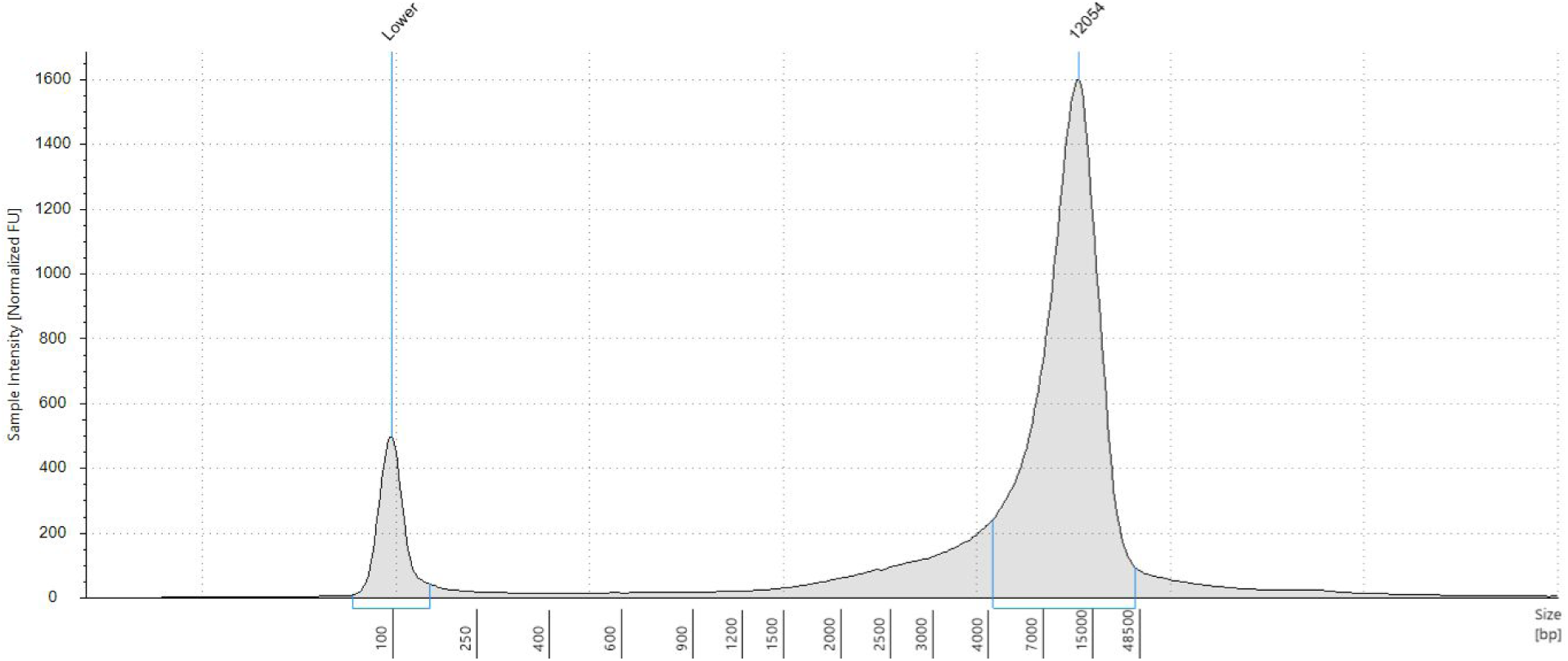
*Tubastraea coccinea.* library size evaluation. Electropherogram traces of the constructed SMRTbell library.

#### *Tubastraea* sp

As for the SMART bell library preparation, 10 µg of total DNA from *Tubastraea* sp. were processed without further shearing the DNA. After the steps to repair the DNA repair and to ligate adapters and DNA that didn’t have SMRTbell primers attached were removed. The average size of the library was 14 kb (Supplementary Figure 16).

**Supplementary Figure 16:**
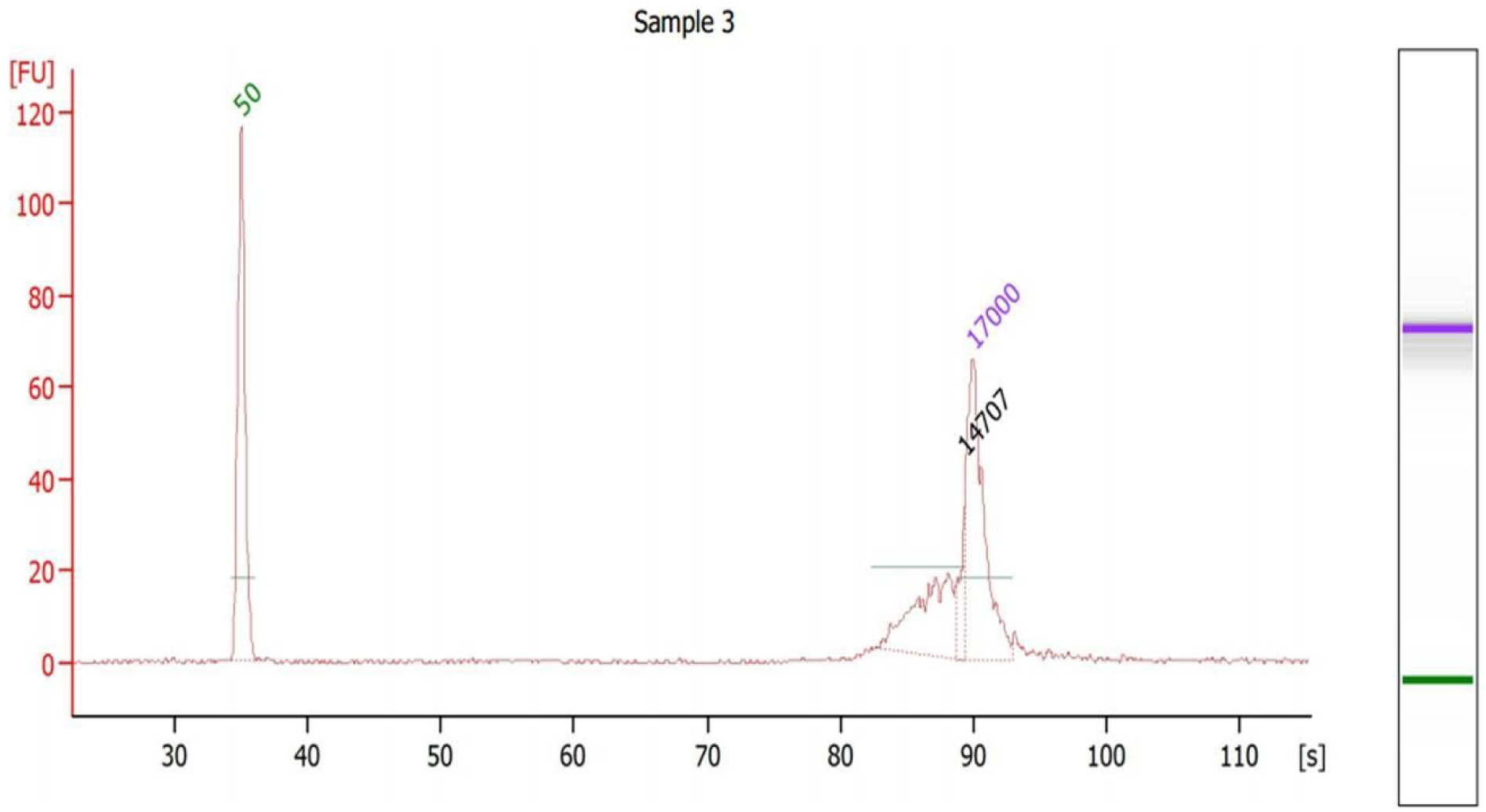
*Tubastraea* sp. library size evaluation. Electropherogram traces of the constructed SMRTbell library.

### 7. Genome assembly and annotation

#### Genome Quality control

The raw data from both experiments were examined using fastQC v.0.11.8 ^60^ and K-mer analysis v.2.4.1 (KAT; ^61^. Long-reads data were also examined using SMRT link analysis v.6.0.0.47836 and stsPlots.

After visual inspection of the fastQC and KAT results, for short-reads we proceeded to the quality control trimming and filtering using bbDUK v.38.42. Illumina PE data was modulo trimmed (ftm = 5) due to the lower quality of the last base in the 151 bp reads. In the second round, reads were quality trimmed using the following parameters: i) minlength=100 (minimum length); ii) ktrim=r (trim reads matching reference k-mers); iii) k=23 (k-mer size used to find contaminants); iv) hdist=1 (Hamming distance for reference k-mers); v) tbo=t (trim adapters based on reads overlap); vi) tpe=t (trimming on both reads); vii) qtrim=rl (trim both reads ends based on quality score); viii) trimq=20 (quality trimming score); and ix) minavgquality=20 (minimum average read quality). Read contaminants were searched against the phix control library and human sequences.

#### Hybrid Assembly

The genome assembly of *Tubastraea* sp. was performed with MaSuRCA v.3.3.1 (Maryland Super-Read Celera Assembler ^62^ using Flye as final assembler and default parameters apart from JF_SIZE = 30,000,000,000. PE was set specifically for each species: i) *Tubastraea tagusensis* = 494 48; ii) *Tubastraea coccinea* = 500 50 ; and iii) *Tubastraea* sp. = 498 50. The quality control of *Tubastraea* spp. assembly was accessed by QUAST v5.0.2 ^63^, BUSCO v3.0.2 ^64^ Benchmarking Universal Single-Copy Orthologs) and stats.sh from BBMap v.38.42. The inflated estimated genome size and high percentage of duplicated gene copies on BUSCO suggested the presence of diploid sequences in the first draft assemblies. Then, to achieve a haploid assembly we used the purge_haplotigs pipeline v1.0.4 ^65^ to remove haplotigs from the draft assemblies. The final statistics from our haploid draft assembly were then compared to other coral genomes retrieved from NCBI’s Genome portal (https://www.ncbi.nlm.nih.gov/genome) (Table 1).

#### Genome Annotation

Following the quality control steps, we proceeded to the annotation of the curated *Tubastraea* spp. genome. Intrinsic repetitive elements were inferred using ab-initio and homology-based approaches by RepeatModeler v1.0.11 and RepeatMasker v4.0.9 (Dfam v3.0, RepBase v20170127, and custom libraries; -gccalc -noisy -xm -xsmall -gff) (Supplementary Table 4). Approximately 50% of bases were masked for *T. tagusensis* and nearly 58% for *T. coccinea* and *Tubastraea* sp. The proportion of repetitive elements in *Tubastraea* spp. is similar to other Cnidaria, invertebrates and birds, albeit lower than those observed in reptiles and mammals.

**Supplementary Table 4:** Summary of repeat elements found in *Tubastraea* spp. genomes.

| Families of repeat elements | Species | Number of elements | Length occupied (bp) | % of sequence |
| --- | --- | --- | --- | --- |
| SINEs | *T. tagusensis* | 116,662 | 18,789,036 | 1.33 |
|  | *T. coccinea* | 63,625 | 10,331,995 | 1.28 |
|  | *Tubastraea* sp. | 51,786 | 7,632,315 | 0.96 |
| LINEs | *T. tagusensis* | 219,798 | 69,895,505 | 4.96 |
|  | *T. coccinea* | 167,423 | 53,031,862 | 6.57 |
|  | *Tubastraea* sp. | 146,325 | 46,955,704 | 5.90 |
| LTR elements | *T. tagusensis* | 72,939 | 25,710,204 | 1.82 |
|  | *T. coccinea* | 48,311 | 21,665,510 | 2.68 |
|  | *Tubastraea* sp. | 40,534 | 21,457,462 | 2.70 |
| DNA elements | *T. tagusensis* | 594,954 | 171,039,953 | 12.13 |
|  | *T. coccinea* | 374,713 | 102,375,514 | 12.68 |
|  | *Tubastraea* sp. | 363,878 | 117,314,854 | 14.75 |
| Unclassified | *T. tagusensis* | 1,990,951 | 402,675,152 | 28.56 |
|  | *T. coccinea* | 1,292,503 | 261,046,415 | 32.34 |
|  | *Tubastraea* sp. | 1,245,311 | 253,581,504 | 31.88 |
| Total interspersed repeats | *T. tagusensis* | 20,875 | 688,109,850 | 48.80 |
|  | *T. coccinea* |  | 448,451,296 | 55.55 |
|  | *Tubastraea* sp. |  | 446,941,839 | 56.18 |
| Small RNA | *T. tagusensis* | 15,036 | 3,999,204 | 0.28 |
|  | *T. coccinea* | 23,630 | 4,596,934 | 0.57 |
|  | *Tubastraea* sp. | 27,237 | 4,402,688 | 0.55 |
| Satellites | *T. tagusensis* | 295,255 | 3,901,714 | 0.28 |
|  | *T. coccinea* | 1,288 | 410,658 | 0.05 |
|  | *Tubastraea* sp. | 2,078 | 611,409 | 0.08 |
| Simple repeats | *T. tagusensis* | 34,053 | 16,180,230 | 1.15 |
|  | *T. coccinea* | 159,621 | 9,169,284 | 1.14 |
|  | *Tubastraea* sp. | 146,849 | 7,995,587 | 1.01 |
| Low complexity | *T. tagusensis* | 34,053 | 1,781,691 | 0.13 |
|  | *T. coccinea* | 16,863 | 889,912 | 0.11 |
|  | *Tubastraea* sp. | 16,013 | 842,895 | 0.11 |

The soft-masked genome was annotated using BRAKER v2.1.3 (--cores 32 --crf --gff3 --softmasking --AUGUSTUS_ab_initio --UTR=on). BRAKER encompasses different tools to train, predict and annotate gene structures in an automated fashion. In this study, we relied on hints provided by the alignment of RNA reads to the genome using STAR v2.7.2a. GeneMark-ET, Augustus ab-initio and CRF were used to generate a set of training genes for

Augustus. For gene prediction, we used BRAKER v2.1.3 with the aligned the RNA-seq bam file generated by STAR v.2.7.2a as support for gene models (--softmasking --AUGUSTUS_ab_initio --crf --UTR=on --gff3). At first, hints were generated from the bam file and then processed to feed GENEMARK-ET. The gene models obtained were then filtered to be used in Augustus training. The CRF predictions performed worse than HMM and were discarded in favor of the latter. BRAKER also used Augustus for ab-initio and UTR predictions to refine gene models. The final set of genes models consisted of 10.6% of the genome covered by CDS. The summary statistics for gene models predicted in the genome are shown in Supplementary Table 5. The functional annotation of predicted genes was performed using InterProScan v5.36-75.0 with Pfam v32.0 and PANTHER v14.1 as databases. The predicted genes with homology evidence were filtered to retrieve metabolic pathways and Gene Ontology annotations from PANTHER website using Panther generic mapping identifiers. Whole-genome orthologous gene comparisons and annotations across six Cnidaria species, *Nematostella vectensis, Acropora digitifera, Orbicella faveolata, Tubastraea coccinea, Tubastraea tagusenis* and *Tubastraea* sp. were done using OrthoVenn2 web platform ^66^.

